# Molecular insight into the mechanism of action of some beneficial flavonoids for the treatment of Parkinson’s Disease

**DOI:** 10.1101/2023.04.21.537830

**Authors:** Sima Biswas, Anindita Mitra, Sreekanya Roy, Rita Ghosh, Angshuman Bagchi

## Abstract

Parkinson’s Disease (PD) is a severe neurodegenerative disorder. It is characterized by the declination of dopaminergic (DA) neurons of substantia nigra pars compacta in the mid brain. Decrease in dopamine level in substantia nigra is a major event in PD. Mutation in Parkin protein is responsible for the onset of Autosomal Recessive Juvenile Parkinson’s (ARJP) Disease. Till date, there is no available treatment of PD without any side-effects. Therefore, the focus has been shifted to identifying natural product inhibitors for the treatment of the disorder. Flavonoids, a class of natural products, have proven neuroprotective effects. Some of the flavonoids have their abilities to influence the activities of central nervous system. Therefore, many studies are being conducted to analyze their activities to lower the progression of neurodegenerative diseases. This study was conducted to identify the flavonoids to be used as potential drugs for PD. From the literature, we picked up the flavonoids active on the nervous systems of human beings. We employed a literature mining approach to build a Structure Activity Relationship (SAR) to measure their efficacies. We performed molecular docking simulations using the flavonoids as the ligands and computed their binding free energy values. Our study would therefore point towards future drug-designing endeavours to come up with plausible therapeutics against PD onset.

## INTRODUCTION

In 1817, Parkinson’s Disease (PD), one of the most severe neurodegenerative disorders, was described by James Parkinson in his monograph entitled “the Shaking Palsy” Parkinson, 2002). The age-dependent disorder PD is currently affecting approximately 1% of the population worldwide (DeMaagd & Philip, 2015). Substantia nigra pars compacta’s dopaminergic (DA) neurons are progressively lost in PD thereby affecting the dopamine levels in the brain of the patients (Alexander, 2004) (Triarhou, 2013) (Song & Kim, 2016). Decrease in the dopamine level and accumulation of Lewy bodies into the brain cells lead to various disabling motor symptoms like rigidity, postural instability, bradykinesia and resting tremor. Along with the motor symptoms, numerous non-motor symptoms such as anxiety, hallucinations, depression, and cognitive impairment are also observed and consequently decrease the quality of life of the affected patients. Aggregation and accumulation of cytoplasmic inclusion bodies like α-synuclein, ubiquitin, neurofilaments, and different ubiquitinated proteins are the other hallmarks of PD pathogenesis (Chung et al., 2001) (Kouli et al., 2018) (Gómez-Benito et al., 2020). In most cases, age plays an important role as a potential risk factor for the onset of PD (Hindle et al., 2010) (Collier et al., 2011). Although PD is an age-related disorder, the combined effects of various environmental factors, like exposures to pesticides and herbicides, toxic substances, and heavy metals as well as different genetic factors also lead to the development of PD (Ball et al., 2019) (Pang et al., 2019). Furthermore, rural living, well-water drinking, and farming activities too instigate the process of PD onset (Gatto et al., 2009) (Chin-Chan et al., 2015) (Cagac, 2020). Initially, PD is categorized into two forms: sporadic and familial (Spataro et al, 2015) (Chai & Lim, 2013). Many gene mutations are reported as the causative agents of familial form of PD; among them, the associations of only six of them have so far been explored and confirmed. α- synuclein (Stefanis, 2012) and LRRK2 (Li et al., 2014) gene mutations lead to the onset of autosomal dominant form of PD. On the other hand, Parkin, PINK1, DJ1, and ATP13A2 (Yang & Xu, 2014) gene mutations are associated with the development of autosomal recessive PD (Cookson, 2012) (Nuytemans et al, 2010) (Guo et al., 2008). DJ1 gene mutations are responsible to cause sporadic as well as familial forms of PD. Oxidative stress, mitochondrial dysfunction, and neuro-inflammation are involved in PD pathophysiology(Blesa et al., 2015) (Picca et al., 2020). In contrast, it is found in many epidemiological studies that smoking as well as regular caffeine and tea consumption have some protective effects against PD (Hernán et al, 2002) (Baron, 1996) (Tan et al., 2003) (Ren & Chen, 2020). Since long, scientists are working to determine a suitable therapeutic agent for the treatment. However, most, if not all, of the existing therapeutics have considerable side effects (Zahoor et al, 2018) (Emamzadeh & Surguchov, 2018). Therefore, attempts are being made to determine the natural product inhibitors to treat PD with minimal to no side effects. Phytochemicals are one such group of natural products; they are believed to lower the risk of many neurodegenerative diseases like the Alzheimer’s Disease (AD) and PD (Teles et al., 2018). Phytochemicals are generally obtained from plant sources. Flavonoids, alkaloids, terpenoids and sesquiterpenes are four major groups of phytochemicals (Kumar& Pandey, 2013). In this particular work, we are concerned only with the flavonoid molecules and their protective roles against PD. The flavonoids have proven abilities to act as neuroprotectants among all the phytochemicals (Teles et al., 2018) (Kumar& Pandey, 2013).

### Flavonoids and Parkinson’s Disease

Water-soluble flavonoid molecules are one of the largest families of secondary metabolites from plant sources (Hussein & El-Anssary, 2019) (Yonekura-Sakakibara et al, 2019). Generally, polyphenolic flavonoid molecules are found in various plant-derived foods and beverages, such as grains, green and black tea, onions, hot peppers, kale, broccoli, rutabagas and spinach (Thilakarathna. & Rupasinghe, 2013) (Falcone Ferreyra et al, 2012). Based on their chemical structure flavonoids can be categorized into flavanonol, flavones, isoflavanoids, flavonols, flavanones, chalcones and anthocyanin subgroups (Panche et al., 2016).

In recent years, flavonoid molecules are examined for their therapeutic potentials (Ullah et al., 2020). Surprisingly, in oxidative damage-related diseases such as neurodegenerative diseases (PD and AD), asthma, cancer, and atherosclerosis, flavonoids show potential health benefits to overcome the disease conditions (Atrahimovich et al., 2021) (Adedayo et al., 2015). For this study, we first selected different flavonoid molecules which have therapeutic potentials as neuroprotectants from the literature (Jung & Kim, 2018) (Magalingam et al., 2015) (de Andrade Teles et al., 2018).

We then performed molecular docking simulations of the flavonoids as the ligands and the Parkin protein as the receptor to check their binding efficacies. After that a literature mining approach was employed to further increase the mechanism of action of these flavonoids as neuroprotectants. We used the method of Quantitative Structure Activity Relationship (QSAR) to correlate the binding mechanisms of the flavonoids, measured in terms of their binding free energy values, with their structures. Our work for the first time predicted the plausible mechanistic aspects of the bindings of the flavonoids as putative neuro-therapeutics. Our work is the first step towards a rational drug designing endeavour for the treatment of PD using natural product inhibitors with practically no side-effects.

## MATERIALS AND METHODS

### Structure retrieval and quality assessment

This study targeted the discovery of potential flavonoids against Parkin. So, for this study the crystal structure of the target receptor protein, i.e., Parkin, was required. The structure of the active form of Parkin was available from the Protein Data Bank (PDB) (https://www.rcsb.org/) (Berman et al., 2000). The 3D coordinates of the atoms of the amino acid residues of Parkin protein was downloaded in PDB format using the PBD ID: 4I1H. This structure has some missing residues. So, a full chain model was prepared by inserting the missing amino acid residues using the tool MODELLER (Zimmermann et al., 2018) (Gabler et al., 2020) (Webb & Sali, 2016). Then, the structure was modified by Discovery Studio 2.5 (DS2.5) software suit. The stereo-chemical qualities of the modelled Parkin protein were analyzed using SAVES 6.0 server (https://saves.mbi.ucla.edu/) and Ramachandran plots (Ramachandran & Sasisekharan, 1968) were drawn. All amino acid residues of the protein were found to be present in the allowed regions of the Ramachandran plot indicating the stereo-chemical fitness of the structure. The relationship between amino acid sequence of a protein and the corresponding structure was analyzed by Verify3D (Eisenberg et al., 1997). The ERRAT (Colovos & Yeates, 1993) score was found to be 74.8344% which would also depict a good model quality.

### Retrieval and optimization of Ligand molecules

Apigenin, baicalein, chrysin, daidzein, epicatechin, kaempferol, luteolin, malvidin, morin, naringenin, nobiletin, pelargonidin, quercetin, tangeritin, 6-hydroxyflavone, aromadedrin, azaleatin, biochanin, butin, catechin, cyanidin, peonidin, tricin, wogonin, taxifolin, rhamnetin, rhamnazin, kaempferide, hirsutidin, fustin, diosmetin, myricitrin, fisetin, myricetin, galangin, gossypetin, naringin, hesperetin, eriodictyol, sakuranetin, epigallocatechin, (–)-epigallocatechin-3-gallate (EGCG), theaflavin, phloretin, dihydromorin, garbanzol, dihydrogossypetin, genistein, isorhamnetin, delphinidinare, rutin, 7,8- dihydroxyflavone (7,8-DHF), petunidin, ampelopsin, hesperidin, astilbin, pinocembrin, puerarin, silibinin B and calycosin are the flavonoid molecules, reported to have neuroprotective functions in literatures (Magalingam et al., 2015), (Jung & Kim, 2018). The structures of the molecules in .sdf were obtained from PubChem Database (http://www.pubchem.ncbi.nlm.nih.gov/) along with their physical and chemical properties. The structures so obtained in .sdf were ultimately converted to the necessary .pdb format in DS2.5.

### ADMET and Lipinski Filter analysis of selected candidate flavonoids

We used the server Lipinski Filter (http://www.scfbio-iitd.res.in/software/drug design/lipinski. jsp) to determine the drug likeliness (Supplementary spreadsheet) properties of the aforementioned ligands with the help of Lipinski’s Rule of Five (Lipinski, 2004) which helps to discriminate between a drug and a non-drug like molecule based on the following parameters: Molecular Mass (should be less than 500 Dalton)

High Lipophilicity (expressed as log P with a value less than 5) Presence of less than 5 numbers of hydrogen bond donors Presence of less than 10 numbers of hydrogen bond acceptors and Molar refractivity (should lie in the range 40 to 130).

Success or failure of a molecule to act as a drug depends on their satisfactory fulfilments of these criteria. Any molecule to be considered as a potential drug candidate must possess any two or more of the aforementioned drug likeliness properties. Furthermore, admetSAR (http://lmmd.ecust.edu.cn/admetsar2) (Yang et al., 2019) tool was used to analyze ADMET (adsorption, distribution, metabolism, excretion and toxicity) properties of the ligands. The ADMET properties are used to check the presence of necessary characteristics required for a molecule to act like a candidate drug.

### Prediction of active site regions

To identify the distributions of amino acid residues in the active site of Parkin, we used Castp Server (http://sts.bioe.uic.edu). The structure of Parkin was submitted to the server to determine the ligand binding sites present in the protein. We collected the binding site information of Parkin from literature as well. All of this information was utilized in our docking simulations.

### Preparation of protein and ligand molecules

We used the ‘clean protein’ module in DS2.5 to correct the bond lengths and bond angles in the Parkin protein. The atomic charges were calculated using CHARMm force-field (Brooks et al., 2009). The optimizations of the ligand molecules were performed by MMFF force field, suitable for organic ligands (Halgren et al., 1996), by using prepare ligand protocol of the DS2.5.

### Molecular docking analysis and SAR studies

Molecular docking simulations were carried out using the Autodock4.2 software. The principle of Lamarckian Genetic Algorithm is used in Autodock4.2. Parkin protein was kept rigid throughout the docking simulations whereas the ligands were allowed to move. For each ligand, the tool came up with 10 different poses of the receptor-ligand complexes. The AutoDock4.2 scoring functions ranked and scored the poses based on their binding free energy values. After that, we performed a SAR analysis of the candidate flavonoids by literature mining. This was performed to strengthen our claims regarding the efficacies of the candidate ligands as potential neurotherapeutics.

## STRUCTURE-BASED QUANTITATIVE STRUCTURE ACTIVITY RELATIONSHIP (QSAR) STUDIES

The flavonoids were grouped into anthocyanins, flavanones, flavones, flavonols, isoflavonoids, chalcones, and flavanonol based on their backbone structures. Structure-based quantitative structure activity relationship studies were performed to see for any correlation between the pharmaco-kinetic or physico-chemical or the drug likeliness properties of the flavonoids with their binding efficiencies to the Parkin protein. Different physico-chemical properties associated with the drug likeliness of a molecule were calculated using Lipinski filter & admetSAR tools. We chose the mostly used drug likeliness properties, such as XLOGP3, Molecular Weight (MW), Topological Polar Surface Area (TPSA), Blood Brain Barrier (BBB), and Water Solubility for our analysis. The Autodock binding free energy (B.F.E) values were obtained from docking the flavonoids to the Parkin protein using the tool Autodock4.2. This was done for investigation of any dependence of these pharmacokinetic properties on the binding free energy values (Bellocchi et al., 2005) (Mitra et al., 2019). Firstly, all the pharmacokinetic properties were compared individually with the Autodock B.F.E. for building a linear regression model. Keeping the Autodock B.F.E in the X-axis and one of the aforementioned drug likeliness properties in the Y-axis, any possible correlation between the two parameters was determined by calculating the co-efficient of determination or R^2^ value. We then tested the predictivity of the linear regression model using training-testing validation. Both the datasets i.e. MW of flavonoids and their corresponding Autodock B.F.E values were grouped in different strata according to their backbone structures (like anthocyanins, flavanones, flavones, flavonols, isoflavonoids, chalcones, and flavanonol). Stratified random sampling was then done for each strata using Microsoft excel 2010. Conventional training-testing validation was performed, where we selected 40 flavonoids in the training set and 20 flavonoids in the test set. In order to produce a regression model equation with a higher degree of predictivity, we created our training and testing datasets in such a way that each of them should contain representatives from all seven of the flavonoid groups. The training – test analysis was subjected to three fold cross validation, each time with different combinations of flavonoids obtained from stratified random sampling from Microsoft excel 2010. This was done to avoid sampling biasness. The co-efficient of determination R^2^ was then calculated in testing the validity of the built model equations. The predictivity of the model was also cross-validated using both internal and external cross- validation metrices (Shamsara et al., 2017). External cross-validation of the training-test set analysis was done to verify further the degree of correlation present between the observed MW and predicted MW from the test set molecules using the regression model equation by finding the R_2 pred_ value following Roy et al.,2015 (Roy et al., 2015) using the equation-

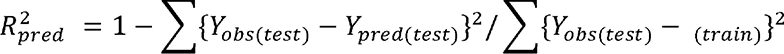

where,

*Y_obs(test)_*: the observed MW of the test set molecules,

*Y_pred(test)_*: the predicted MW for the test set molecules from the regression model equation,

LJ*_train_*: the mean value of the observed MW of the training set molecules.

Cross-validation through Leave-One-Out (LOO) mechanism was also performed and the predicted residual sum of squares (PRESS) and cross-validated R^2^ (Q^2^) were calculated using the following equation as mentioned in (Roy et al., 2015) -

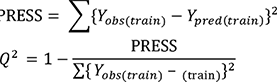

where,

*Y_obs(train)_*: the observed MW of the training set molecules,

*Y_pred(train)_*: the LOO-derived predicted MW of the training set molecules, LJ*_train_*: the mean value of the observed MW of the training set molecules. **RESULTS**

### Analysis of the physicochemical characteristics of Parkin protein

We calculated different physico-chemical characteristics of Parkin protein using the tool ProtParam (Gasteiger et al., 2005). The computed molecular weight and isoelectric point of Parkin protein were found to be 36393.53 Da and 7.06 respectively. The Grand Average of Hydropathicity (GRAVY) and aliphatic index were found to be -0.372 and 60.28 respectively. The negative GRAVY value of Parkin protein would indicate that the protein is non-polar. The high value of the aliphatic index would also point towards the overall non-polar nature of the protein. The instability index (II) of Parkin was computed to be 43.09 indicating that the protein is unstable.

### Structural validation of Parkin protein

The reliability of the 3D structure of the modelled Parkin protein was checked by Ramachandran plot (Supplementary Figure1) where all the amino acid residues were present in the appropriate regions indicating the stereochemical fitness of the protein structure. Further validations of the stereochemical fitness of the protein were performed using Verify3D and ERRAT. The results from these tools established the suitability of the protein model to be used for further analysis.

### Analyses of the drug-likeliness features of the flavonoids

We browsed literature and gathered information about the flavonoids which have proven roles in the prevention of neurodegenerative diseases (Magalingam et al., 2015) (Jung & Kim, 2018). We used Lipiniski’s rule of five to check the drug likeliness properties of the flavonoids (Supplementary Table1). We also checked the ADMET profiles of the ligands to verify their suitability as plausible drug candidates (Supplementary Table2). From the ADMET profile analysis, it was observed that most of the flavonoids have all the properties necessary to act as potential drug candidates. It is to be noted that lipophilicity too serves as an important parameter for a ligand to act as a potential drug candidate for neurological disorders (Alavijeh et al., 2005). Therefore, these ligands can be potential candidates even if they are unable to cross the Blood Brain Barrier (BBB).

### Active site prediction and molecular docking analysis of Parkin protein with ligands

Ligand binding sites on Parkin protein were predicted by CastP server using its 3D coordinates of the atoms of the amino acids. This information of binding site was used to carry out the docking studies. In this case, we used rigid body directed docking.

The amino acid residues present in the binding site of Parkin protein are presented below:

TYR143, THR231, ASN232, SER233, ARG234, ASN235, ILE236, THR237, THR240, ARG256, ARG396, ALA398, GLN400, ARG402, TRP403, GLU404, ALA405, ALA406, SER407, LYS408, GLU409, THR410, LYS412, LYS413, THR414, LYS416, LYS427, GLY430, CYS431, MET432, HIS433, GLU444, TRP447, ASN448, PHE463, ASP464, VAL465 Among the 10 docking poses for each of the Parkin-flavonoid complexes, the best ones were selected based on their binding free energy values. The Parkin-flavonoid complexes with the maximum negative binding free energy values were considered for further analysis (Table1). The binding interactions are presented in Supplementary_flavonoids.pptx.

**Table 1.**
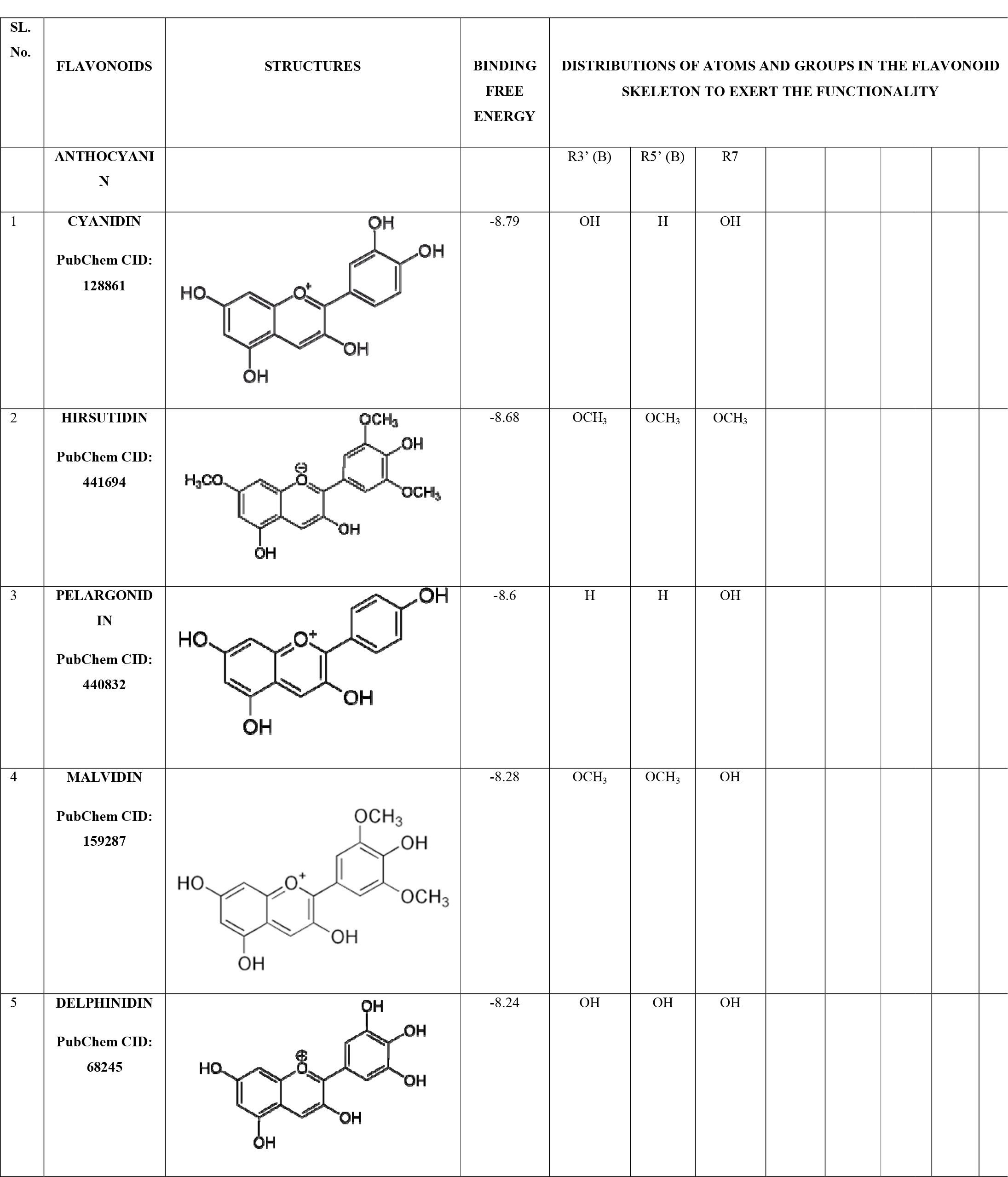
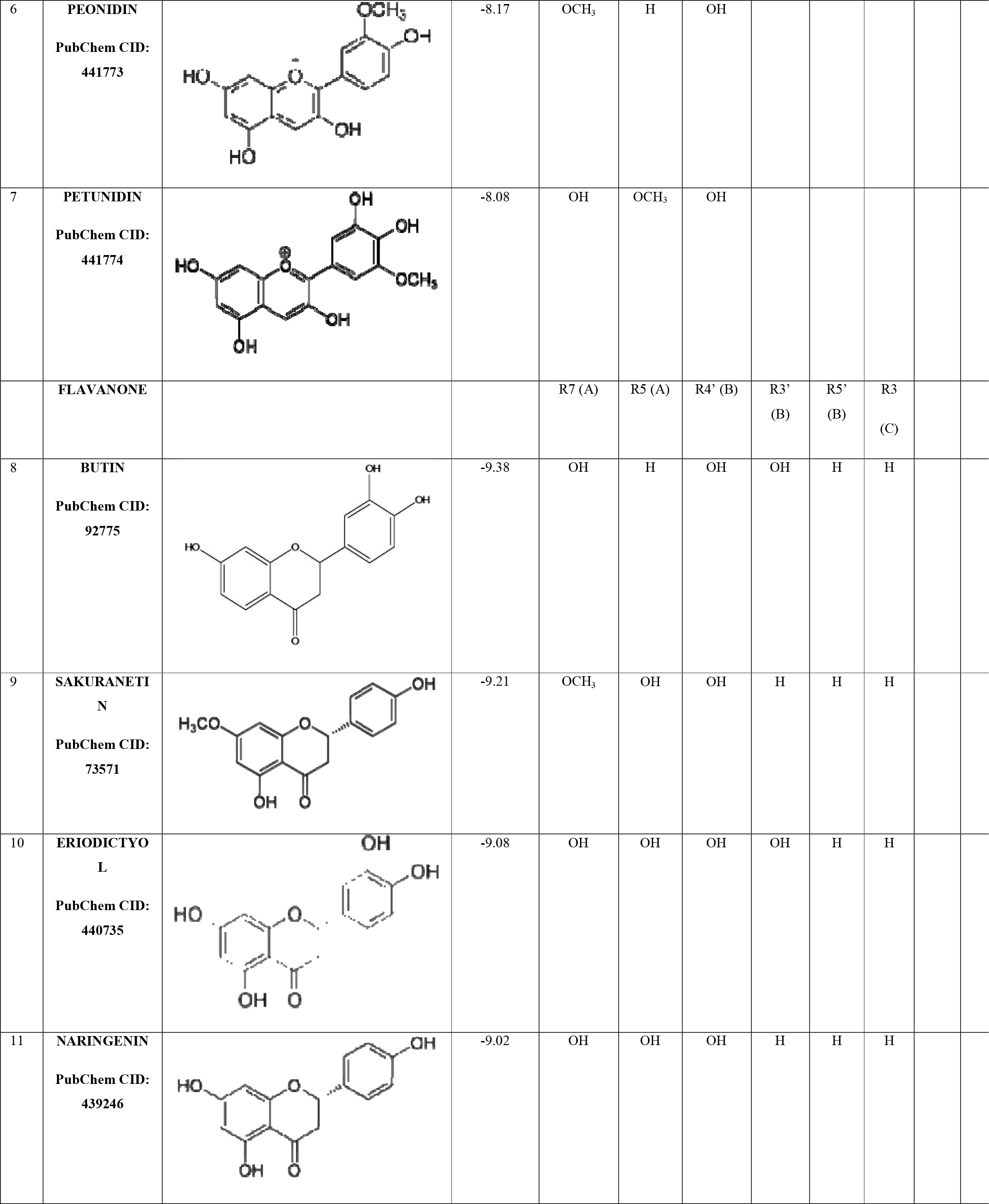
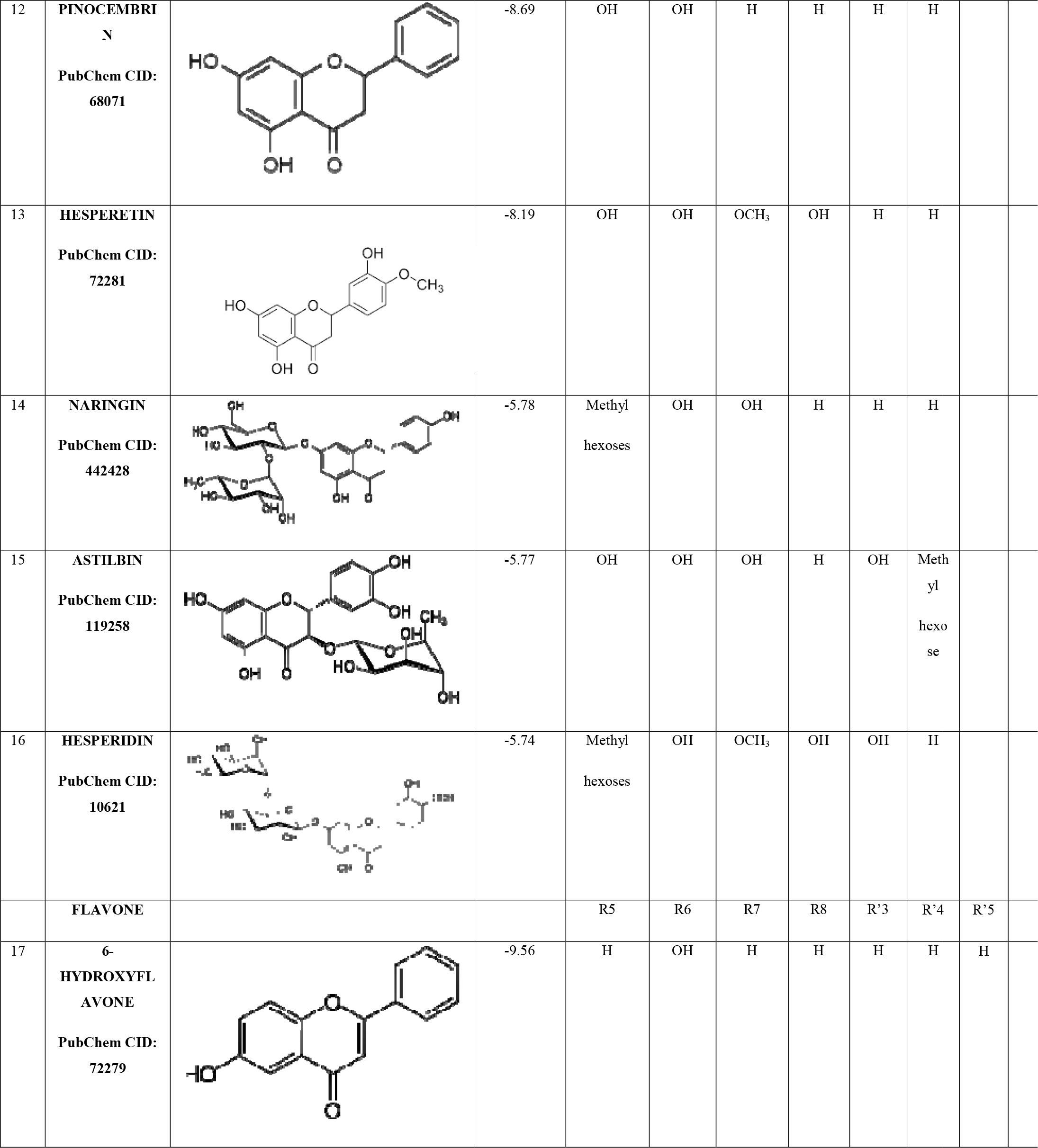
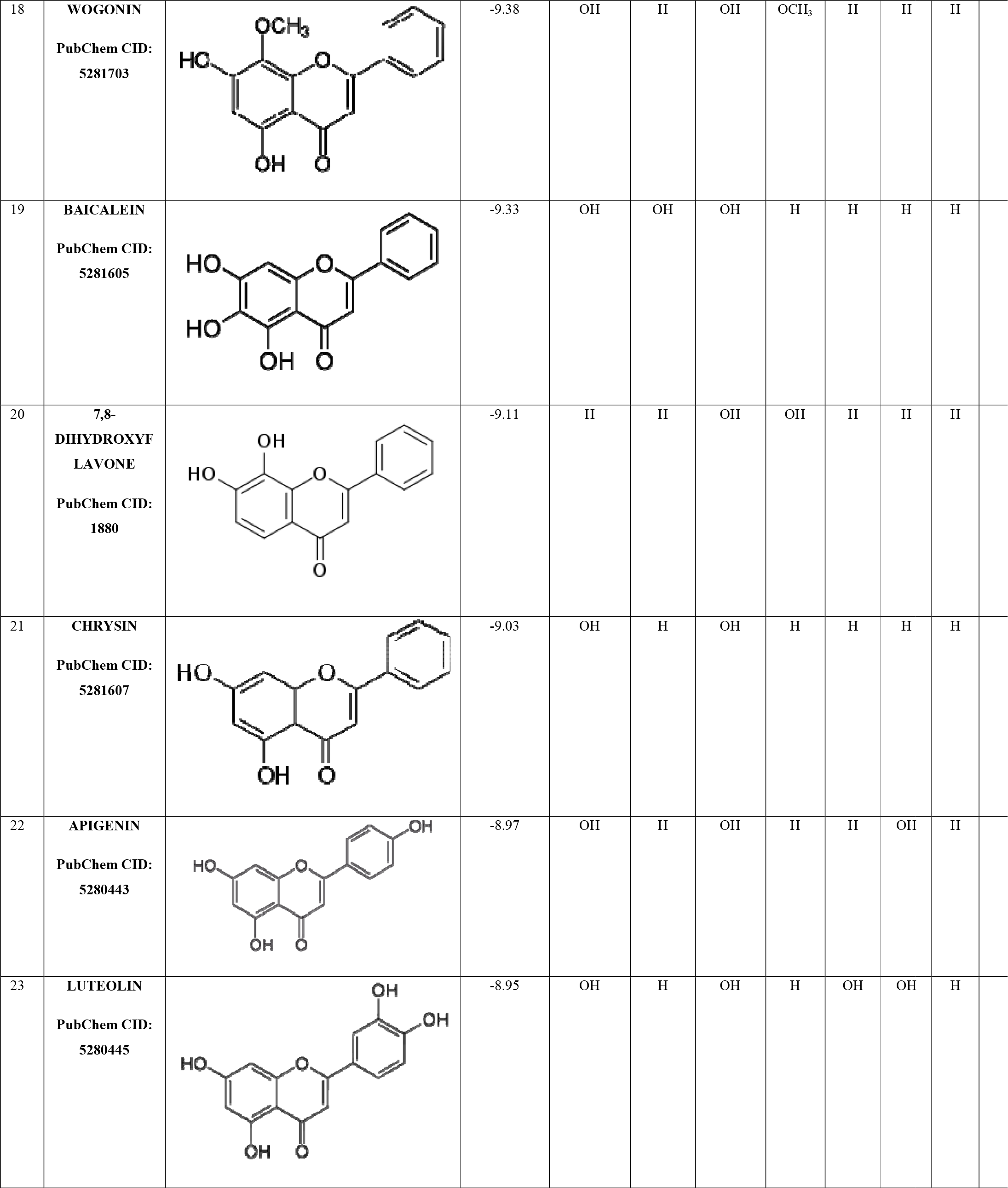
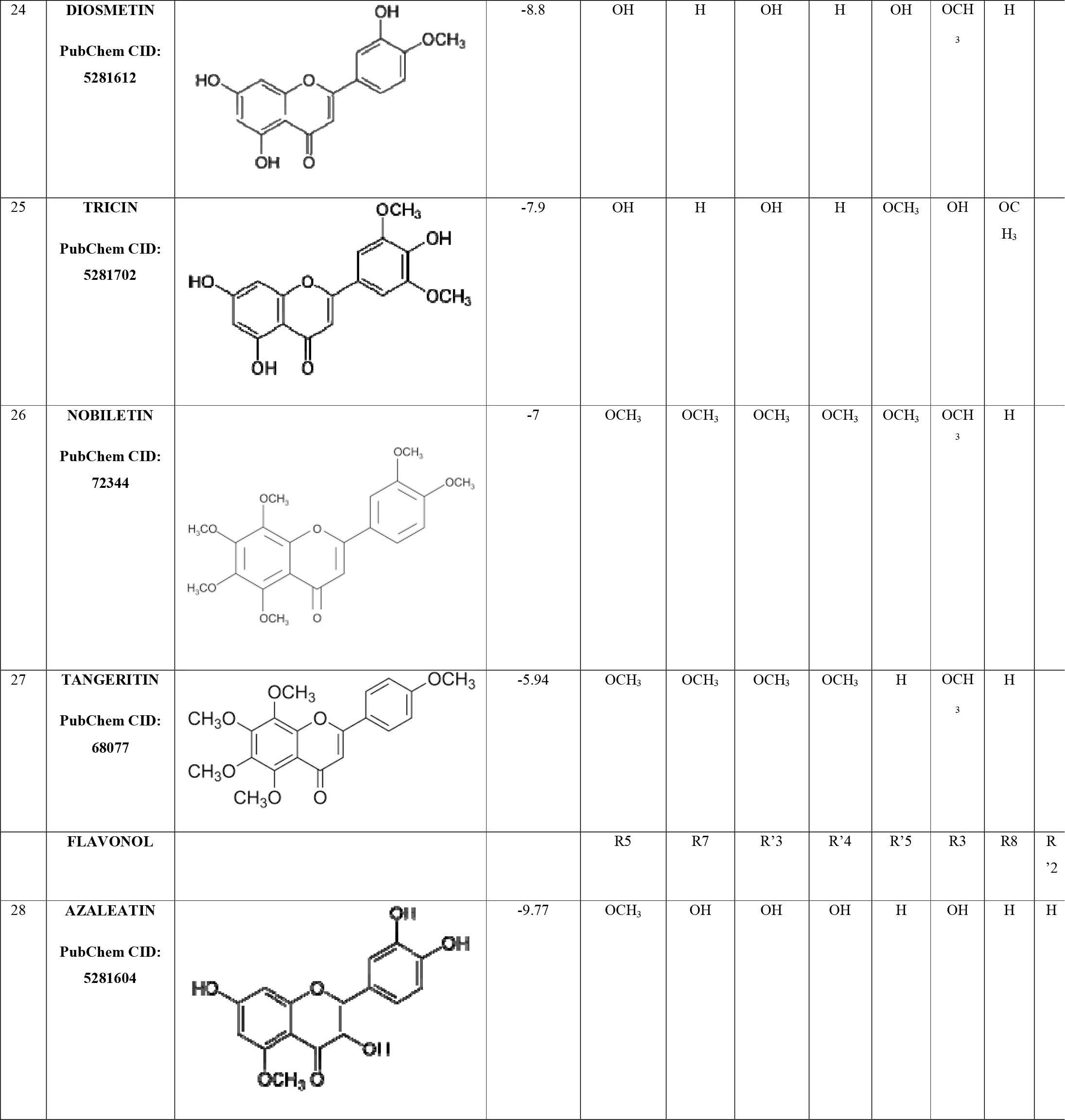
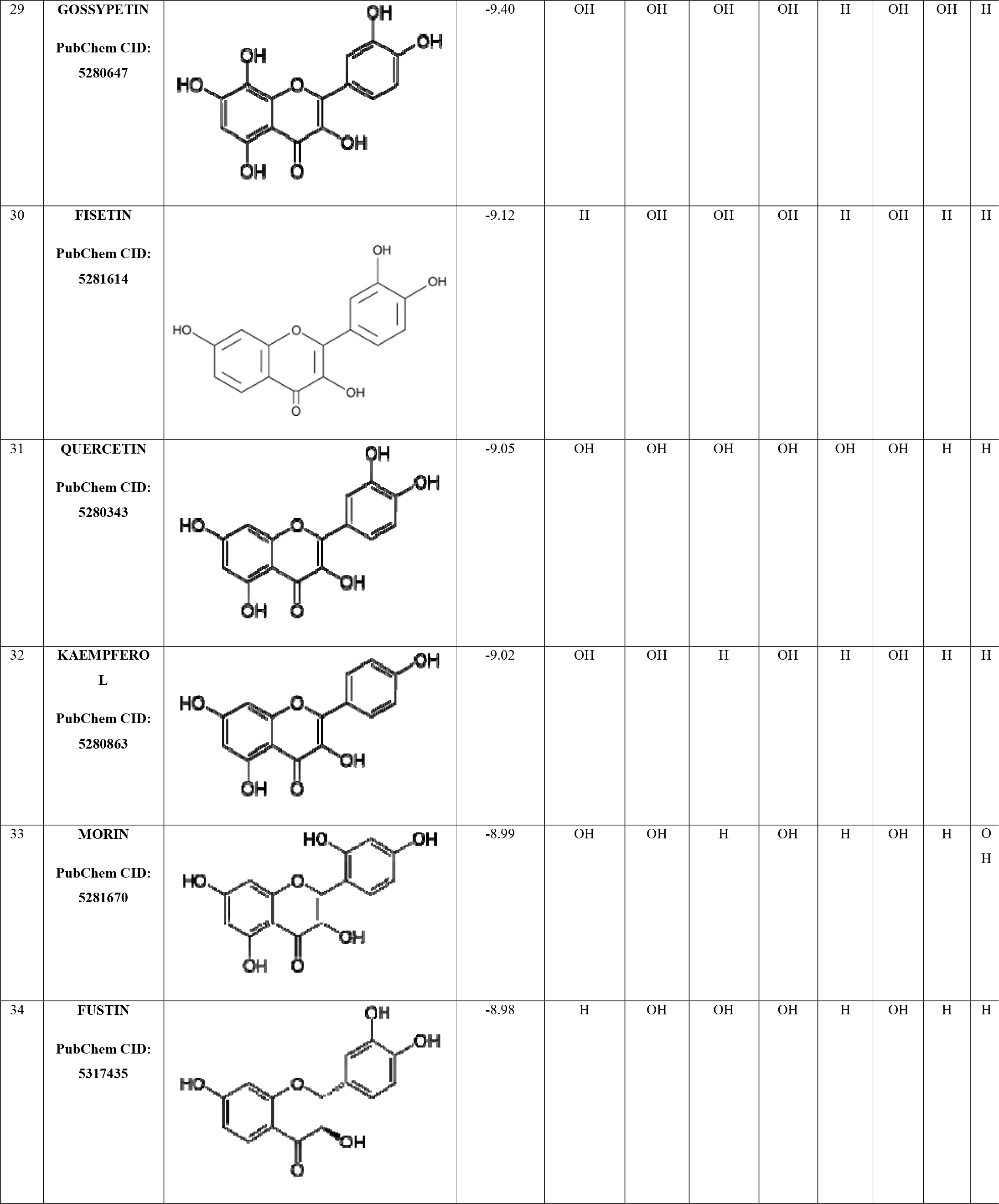
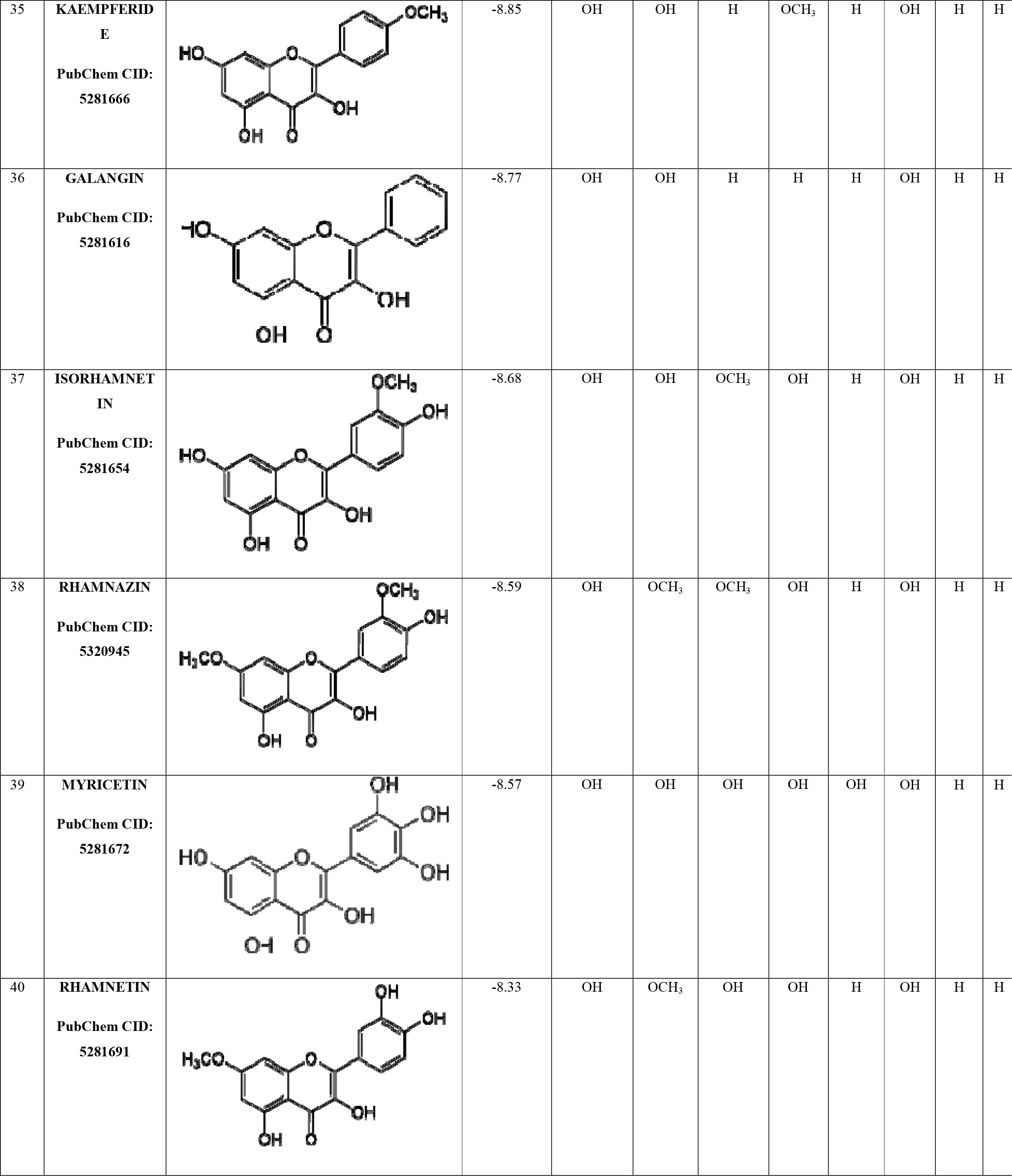
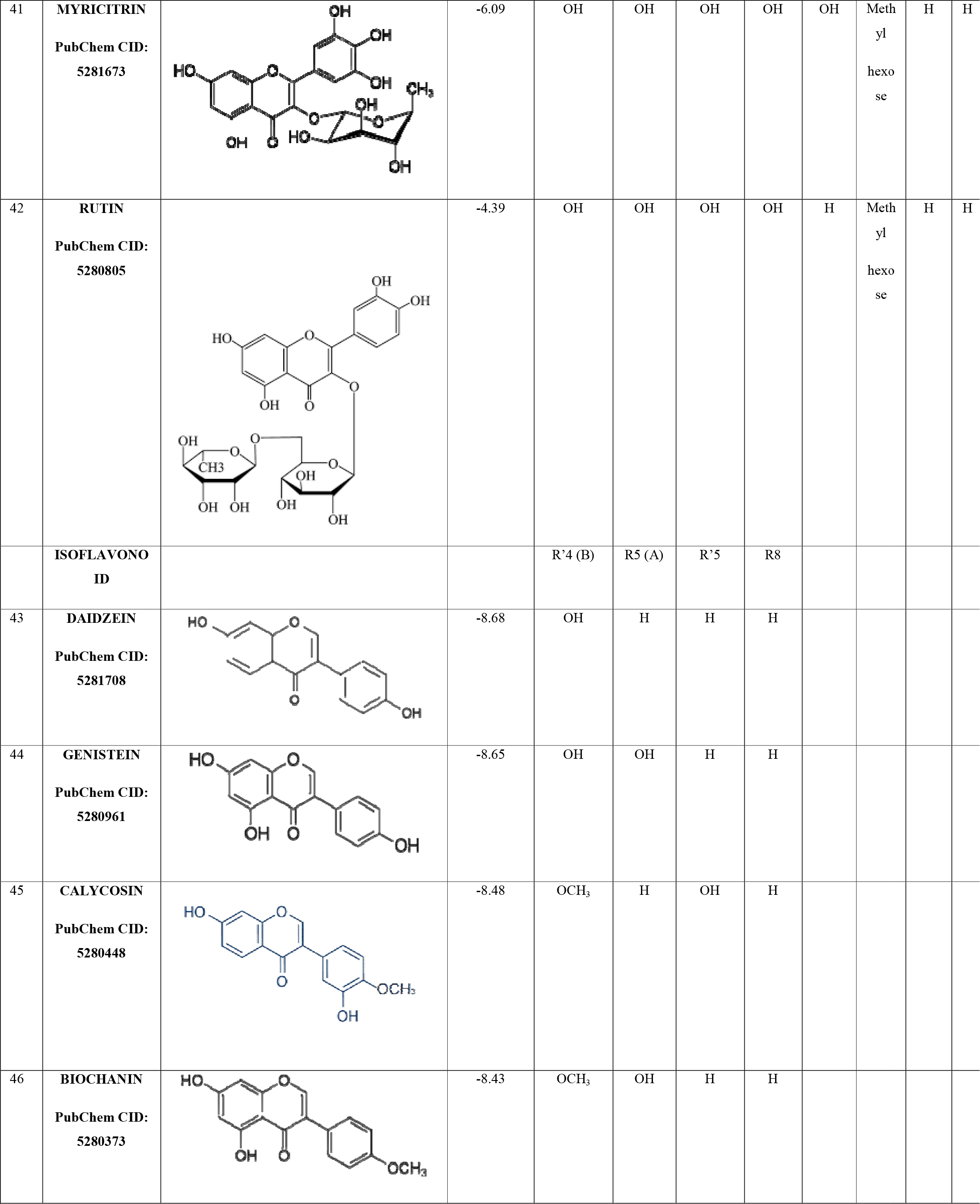
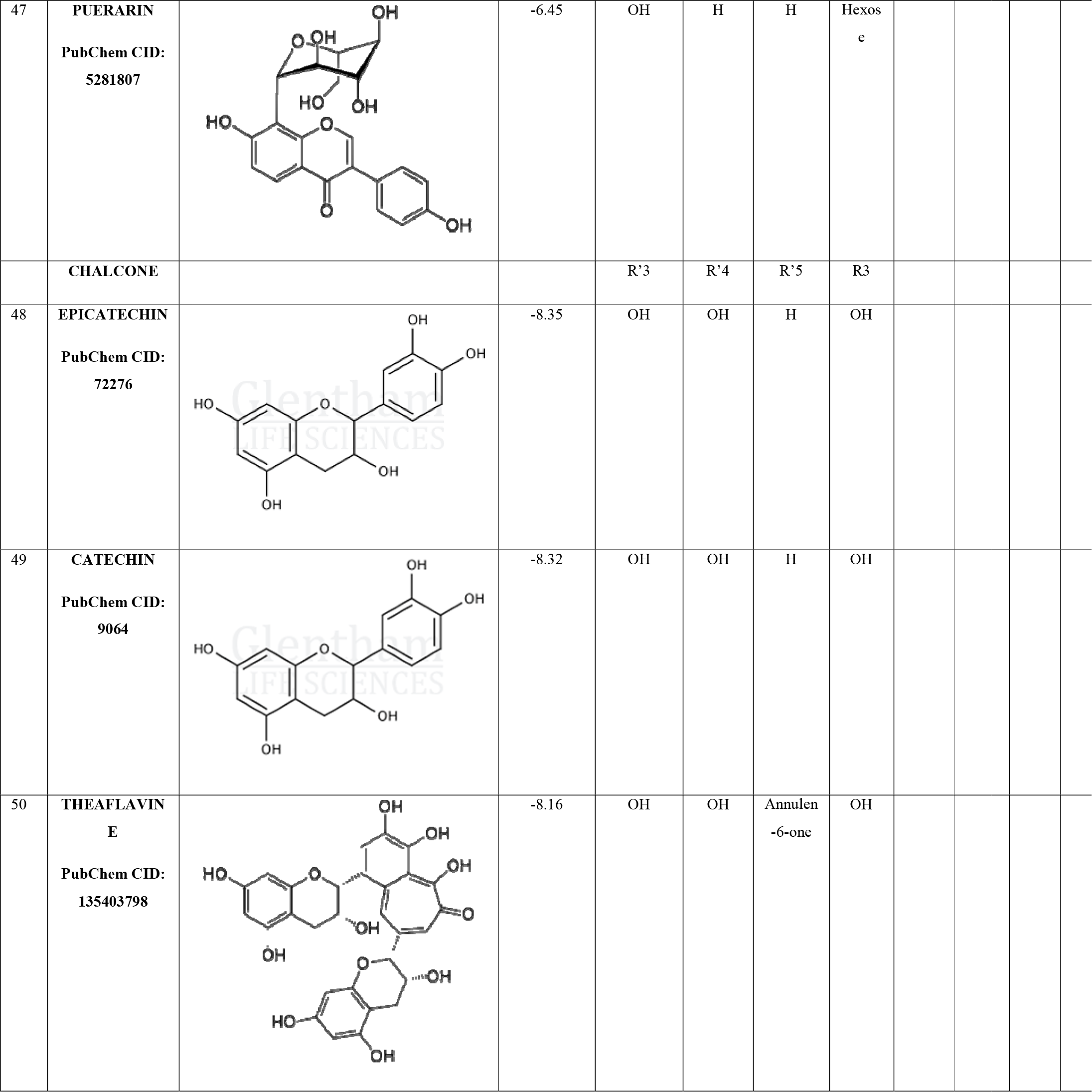
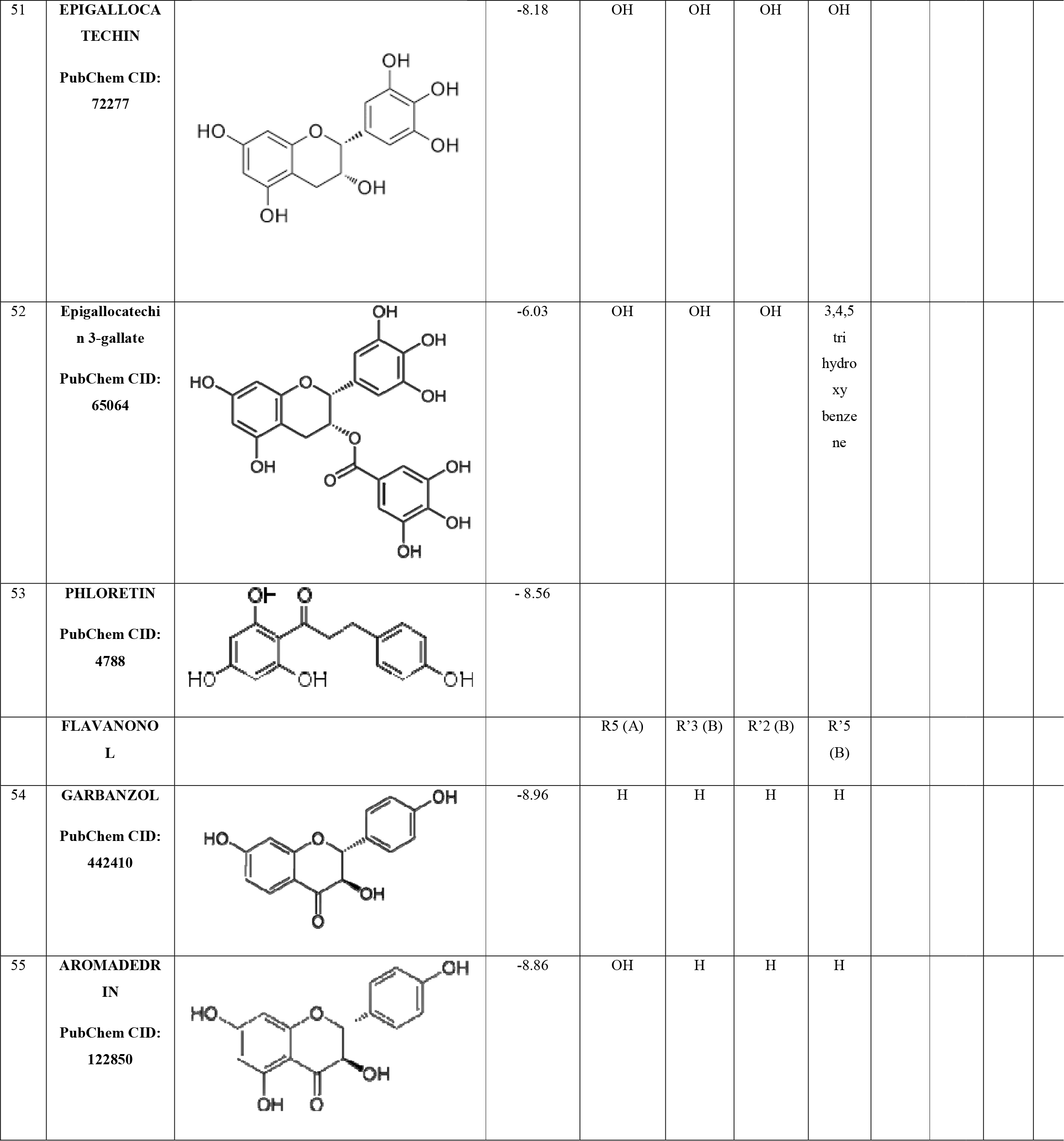
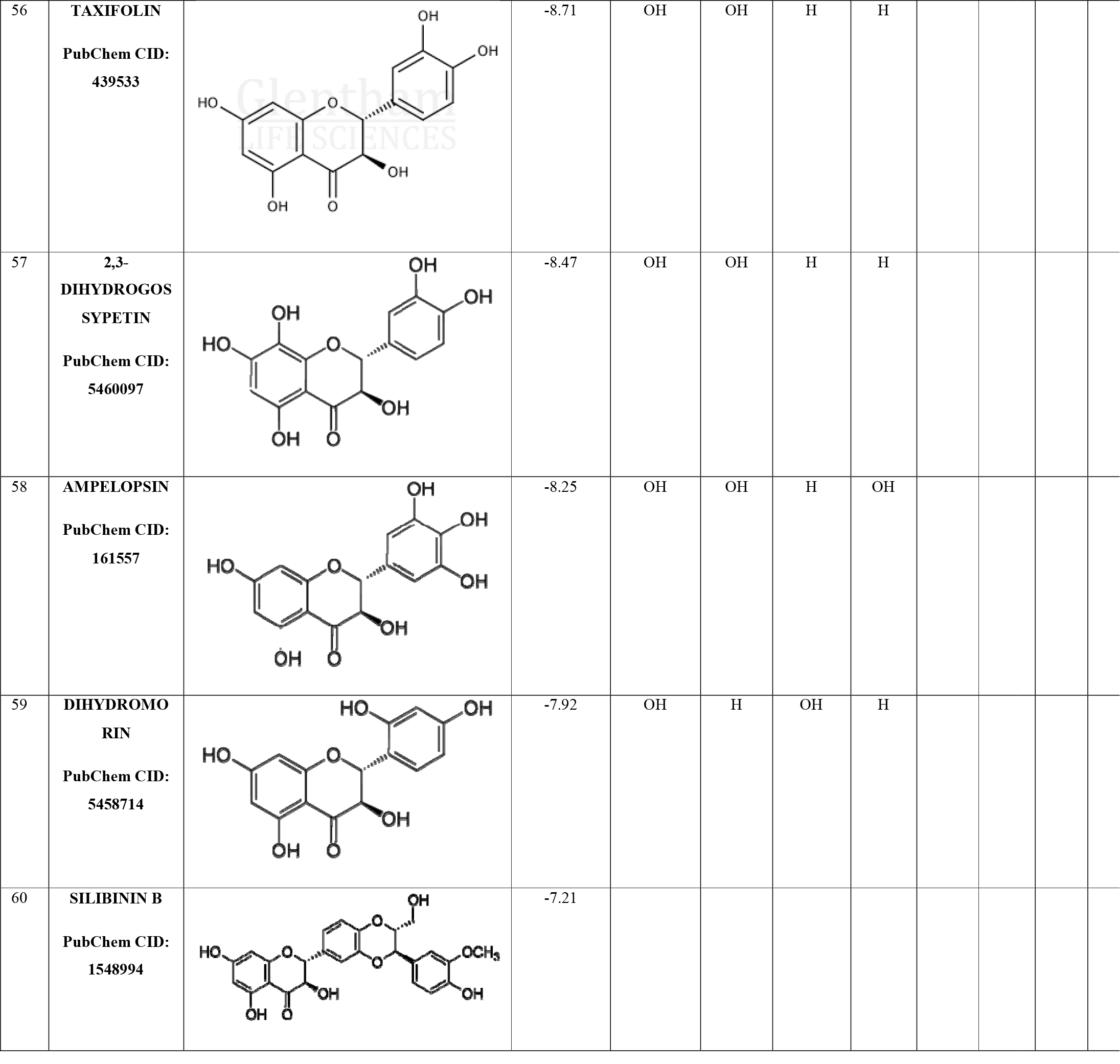
: Binding free energy values of the selected flavonoids interacting with Parkin protein.

| SL. No. | FLAVONOIDS | STRUCTURES | BINDING FREE ENERGY | DISTRIBUTIONS OF ATOMS AND GROUPS IN THE FLAVONOID SKELETON TO EXERT THE FUNCTIONALITY |  |  |
| --- | --- | --- | --- | --- | --- | --- |
|  |  |  |  | R3' (B) | R5' (B) | R7 |
| 1 | **CYANIDIN**<br>PubChem CID: 128861 |  | -8.79 | OH | H | OH |
| 2 | **HIRSUTIDIN**<br>PubChem CID: 441694 |  | -8.68 | OCH<sub>3</sub> | OCH<sub>3</sub> | OCH<sub>3</sub> |
| 3 | **PELARGONIDIN**<br>PubChem CID: 440832 |  | -8.6 | H | H | OH |
| 4 | **MALVIDIN**<br>PubChem CID: 159287 |  | -8.28 | OCH<sub>3</sub> | OCH<sub>3</sub> | OH |
| 5 | **DELPHINIDIN**<br>PubChem CID: 68245 |  | -8.24 | OH | OH | OH |

|  |  |  |  |  |  |  |  |  |  |
| --- | --- | --- | --- | --- | --- | --- | --- | --- | --- |
| 6 | **PEONIDIN**<br>**PubChem CID:**<br>**441773** |  | -8.17 | OCH<sub>3</sub> | H | OH |  |  |  |
| 7 | **PETUNIDIN**<br>**PubChem CID:**<br>**441774** |  | -8.08 | OH | OCH<sub>3</sub> | OH |  |  |  |
|  | **FLAVANONE** |  |  | R7 (A) | R5 (A) | R4' (B) | R3' (B) | R5' (B) | R3 (C) |
| 8 | **BUTIN**<br>**PubChem CID:**<br>**92775** |  | -9.38 | OH | H | OH | OH | H | H |
| 9 | **SAKURANETIN**<br>**PubChem CID:**<br>**73571** |  | -9.21 | OCH<sub>3</sub> | OH | OH | H | H | H |
| 10 | **ERIODICTYOL**<br>**PubChem CID:**<br>**440735** |  | -9.08 | OH | OH | OH | OH | H | H |
| 11 | **NARINGENIN**<br>**PubChem CID:**<br>**439246** |  | -9.02 | OH | OH | OH | H | H | H |

|  |  |  |  |  |  |  |  |  |  |  |
| --- | --- | --- | --- | --- | --- | --- | --- | --- | --- | --- |
| 12 | **PINOCEMBRI**<br>**N**<br>**PubChem CID:**<br>**68071** |  | -8.69 | OH | OH | H | H | H | H |  |
| 13 | **HESPERETIN**<br>**PubChem CID:**<br>**72281** |  | -8.19 | OH | OH | OCH<sub>3</sub> | OH | H | H |  |
| 14 | **NARINGIN**<br>**PubChem CID:**<br>**442428** |  | -5.78 | Methyl<br>hexoses | OH | OH | H | H | H |  |
| 15 | **ASTILBIN**<br>**PubChem CID:**<br>**119258** |  | -5.77 | OH | OH | OH | H | OH | Methyl<br>hexose |  |
| 16 | **HESPERIDIN**<br>**PubChem CID:**<br>**10621** |  | -5.74 | Methyl<br>hexoses | OH | OCH<sub>3</sub> | OH | OH | H |  |
|  | **FLAVONE** |  |  | R5 | R6 | R7 | R8 | R'3 | R'4 | R'5 |
| 17 | **6-**<br>**HYDROXYFLAVONE**<br>**PubChem CID:**<br>**72279** |  | -9.56 | H | OH | H | H | H | H | H |

|  |  |  |  |  |  |  |  |  |  |  |
| --- | --- | --- | --- | --- | --- | --- | --- | --- | --- | --- |
| 18 | **WOGONIN**<br>PubChem CID:<br>5281703 |  | -9.38 | OH | H | OH | OCH<sub>3</sub> | H | H | H |
| 19 | **BAICALEIN**<br>PubChem CID:<br>5281605 |  | -9.33 | OH | OH | OH | H | H | H | H |
| 20 | **7,8-DIHYDROXYF**<br>**LAVONE**<br>PubChem CID:<br>1880 |  | -9.11 | H | H | OH | OH | H | H | H |
| 21 | **CHRYSIN**<br>PubChem CID:<br>5281607 |  | -9.03 | OH | H | OH | H | H | H | H |
| 22 | **APIGENIN**<br>PubChem CID:<br>5280443 |  | -8.97 | OH | H | OH | H | H | OH | H |
| 23 | **LUTEOLIN**<br>PubChem CID:<br>5280445 |  | -8.95 | OH | H | OH | H | OH | OH | H |

|  |  |  |  |  |  |  |  |  |  |  |  |
| --- | --- | --- | --- | --- | --- | --- | --- | --- | --- | --- | --- |
| 24 | **DIOSMETIN**<br>**PubChem CID:**<br>**5281612** |  | -8.8 | OH | H | OH | H | OH | OCH<sub>3</sub><br>3 | H |  |
| 25 | **TRICIN**<br>**PubChem CID:**<br>**5281702** |  | -7.9 | OH | H | OH | H | OCH<sub>3</sub> | OH | OC<br>H<sub>3</sub> |  |
| 26 | **NOBILETIN**<br>**PubChem CID:**<br>**72344** |  | -7 | OCH<sub>3</sub> | OCH<sub>3</sub> | OCH<sub>3</sub> | OCH<sub>3</sub> | OCH<sub>3</sub> | OCH<sub>3</sub><br>3 | H |  |
| 27 | **TANGERITIN**<br>**PubChem CID:**<br>**68077** |  | -5.94 | OCH<sub>3</sub> | OCH<sub>3</sub> | OCH<sub>3</sub> | OCH<sub>3</sub> | H | OCH<sub>3</sub><br>3 | H |  |
|  | **FLAVONOL** |  |  | R5 | R7 | R'3 | R'4 | R'5 | R3 | R8 | R'<br>2 |
| 28 | **AZALEATIN**<br>**PubChem CID:**<br>**5281604** |  | -9.77 | OCH<sub>3</sub> | OH | OH | OH | H | OH | H | H |

|  |  |  |  |  |  |  |  |  |  |  |  |
| --- | --- | --- | --- | --- | --- | --- | --- | --- | --- | --- | --- |
| 29 | **GOSSYPETIN**<br>**PubChem CID:**<br>5280647 |  | -9.40 | OH | OH | OH | OH | H | OH | OH | H |
| 30 | **Fisetin**<br>**PubChem CID:**<br>5281614 |  | -9.12 | H | OH | OH | OH | H | OH | H | H |
| 31 | **Quercetin**<br>**PubChem CID:**<br>5280343 |  | -9.05 | OH | OH | OH | OH | OH | OH | H | H |
| 32 | **KAEMPFEROL**<br>**PubChem CID:**<br>5280863 |  | -9.02 | OH | OH | H | OH | H | OH | H | H |
| 33 | **MORIN**<br>**PubChem CID:**<br>5281670 |  | -8.99 | OH | OH | H | OH | H | OH | H | O<br>H |
| 34 | **FUSTIN**<br>**PubChem CID:**<br>5317435 |  | -8.98 | H | OH | OH | OH | H | OH | H | H |

|  |  |  |  |  |  |  |  |  |  |  |  |
| --- | --- | --- | --- | --- | --- | --- | --- | --- | --- | --- | --- |
| 35 | **KAEMPFERID E**<br>**PubChem CID:**<br>5281666 |  | -8.85 | OH | OH | H | OCH<sub>3</sub> | H | OH | H | H |
| 36 | **GALANGIN**<br>**PubChem CID:**<br>5281616 |  | -8.77 | OH | OH | H | H | H | OH | H | H |
| 37 | **ISORHAMNET IN**<br>**PubChem CID:**<br>5281654 |  | -8.68 | OH | OH | OCH<sub>3</sub> | OH | H | OH | H | H |
| 38 | **RHAMNAZIN**<br>**PubChem CID:**<br>5320945 |  | -8.59 | OH | OCH<sub>3</sub> | OCH<sub>3</sub> | OH | H | OH | H | H |
| 39 | **MYRICETIN**<br>**PubChem CID:**<br>5281672 |  | -8.57 | OH | OH | OH | OH | OH | OH | H | H |
| 40 | **RHAMNETIN**<br>**PubChem CID:**<br>5281691 |  | -8.33 | OH | OCH<sub>3</sub> | OH | OH | H | OH | H | H |

|  |  |  |  |  |  |  |  |  |  |  |  |
| --- | --- | --- | --- | --- | --- | --- | --- | --- | --- | --- | --- |
| 41 | **MYRICITRIN**<br>PubChem CID:<br>5281673 |  | -6.09 | OH | OH | OH | OH | OH | Methyl<br>hexose | H | H |
| 42 | **RUTIN**<br>PubChem CID:<br>5280805 |  | -4.39 | OH | OH | OH | OH | H | Methyl<br>hexose | H | H |
|  | **ISOFLAVONO**<br>**ID** |  |  | R'4 (B) | R5 (A) | R'5 | R8 |  |  |  |  |
| 43 | **DAIDZEIN**<br>PubChem CID:<br>5281708 |  | -8.68 | OH | H | H | H |  |  |  |  |
| 44 | **GENISTEIN**<br>PubChem CID:<br>5280961 |  | -8.65 | OH | OH | H | H |  |  |  |  |
| 45 | **CALYCOSIN**<br>PubChem CID:<br>5280448 |  | -8.48 | OCH<sub>3</sub> | H | OH | H |  |  |  |  |
| 46 | **BIOCHANIN**<br>PubChem CID:<br>5280373 |  | -8.43 | OCH<sub>3</sub> | OH | H | H |  |  |  |  |

|  |  |  |  |  |  |  |  |
| --- | --- | --- | --- | --- | --- | --- | --- |
| 47 | **PUERARIN**<br>**PubChem CID:**<br>**5281807** |  | -6.45 | OH | H | H | Hexose |
|  | **CHALCONE** |  |  | R'3 | R'4 | R'5 | R3 |
| 48 | **EPICATECHIN**<br>**PubChem CID:**<br>**72276** |  | -8.35 | OH | OH | H | OH |
| 49 | **CATECHIN**<br>**PubChem CID:**<br>**9064** |  | -8.32 | OH | OH | H | OH |
| 50 | **THEAFLAVIN E**<br>**PubChem CID:**<br>**135403798** |  | -8.16 | OH | OH | Annulene-6-one | OH |

|  |  |  |  |  |  |  |  |
| --- | --- | --- | --- | --- | --- | --- | --- |
| 51 | **EPIGALLOCA**<br>**TECHIN**<br>**PubChem CID:**<br>**72277** |  | -8.18 | OH | OH | OH | OH |
| 52 | **Epigallocatechin 3-gallate**<br>**PubChem CID:**<br>**65064** |  | -6.03 | OH | OH | OH | 3,4,5<br>tri<br>hydro<br>xy<br>benze<br>ne |
| 53 | **PHLORETIN**<br>**PubChem CID:**<br>**4788** |  | - 8.56 |  |  |  |  |
|  | **FLAVANONO**<br>**L** |  |  | R5 (A) | R'3 (B) | R'2 (B) | R'5<br>(B) |
| 54 | **GARBANZOL**<br>**PubChem CID:**<br>**442410** |  | -8.96 | H | H | H | H |
| 55 | **AROMADEDRI**<br>**N**<br>**PubChem CID:**<br>**122850** |  | -8.86 | OH | H | H | H |

|  |  |  |  |  |  |  |  |
| --- | --- | --- | --- | --- | --- | --- | --- |
| 56 | **TAXIFOLIN**<br>PubChem CID:<br>439533 |  | -8.71 | OH | OH | H | H |
| 57 | **2,3-DIHYDROGOSYPETIN**<br>PubChem CID:<br>5460097 |  | -8.47 | OH | OH | H | H |
| 58 | **AMPELOPSIN**<br>PubChem CID:<br>161557 |  | -8.25 | OH | OH | H | OH |
| 59 | **DIHYDROMORIN**<br>PubChem CID:<br>5458714 |  | -7.92 | OH | H | OH | H |
| 60 | **SILIBININ B**<br>PubChem CID:<br>1548994 |  | -7.21 |  |  |  |  |

## DISCUSSION

A potential drug candidate has to pass Lipinski’s filter and fulfil all ADMET properties for exerting their actions. Especially in case of neuronal disorders, BBB permeability is a major criterion for a ligand to act as a potential drug (Pardridge, 2012). It is reported in literature that a drug for the treatment of neurodegenerative diseases can be effective if it can cross the BBB. On the other hand, as per the Lipinski’s rule of five, the molecular weight, the number of hydrogen bond donors and acceptors and logP of the potential candidate molecule would predict its suitability as an orally consumable drug. Most of the flavonoids tested have passed these two criteria. In this work, we have calculated the binding interactions between the flavonoids and Parkin protein.

Regression analysis was done between the Autodock B.F.E values of the 60 flavonoids with their different pharmacokinetic properties such as XLOGP3, MOLECULAR WEIGHT, TPSA, BBB, WATER SOLUBILITY, to see for any correlation between them, keeping the the pharmaco-kinetic properties in the y-axis and the. Autodock B.F.E. in the x-axis. A possible correlation was observed in case of MW of the flavonoids with their binding free energies showing a co-efficient of determination value (R^2^) of 0.674 as can be seen in Figure 1. Further validation studies were conducted to test the predictivity of this possible correlation. This correlation was first tested by training set-test set validation. The training-test set validation of the correlation was repeated thrice each time with different sets of flavonoids in the training set and the test set (SET I, SET II and SET III), using stratified random sampling, to avoid biasness. The three combinations of the data are listed in (Supplementary Table 3). The training set-test set analysis produced three model equations as

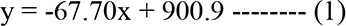

**Figure 1:**
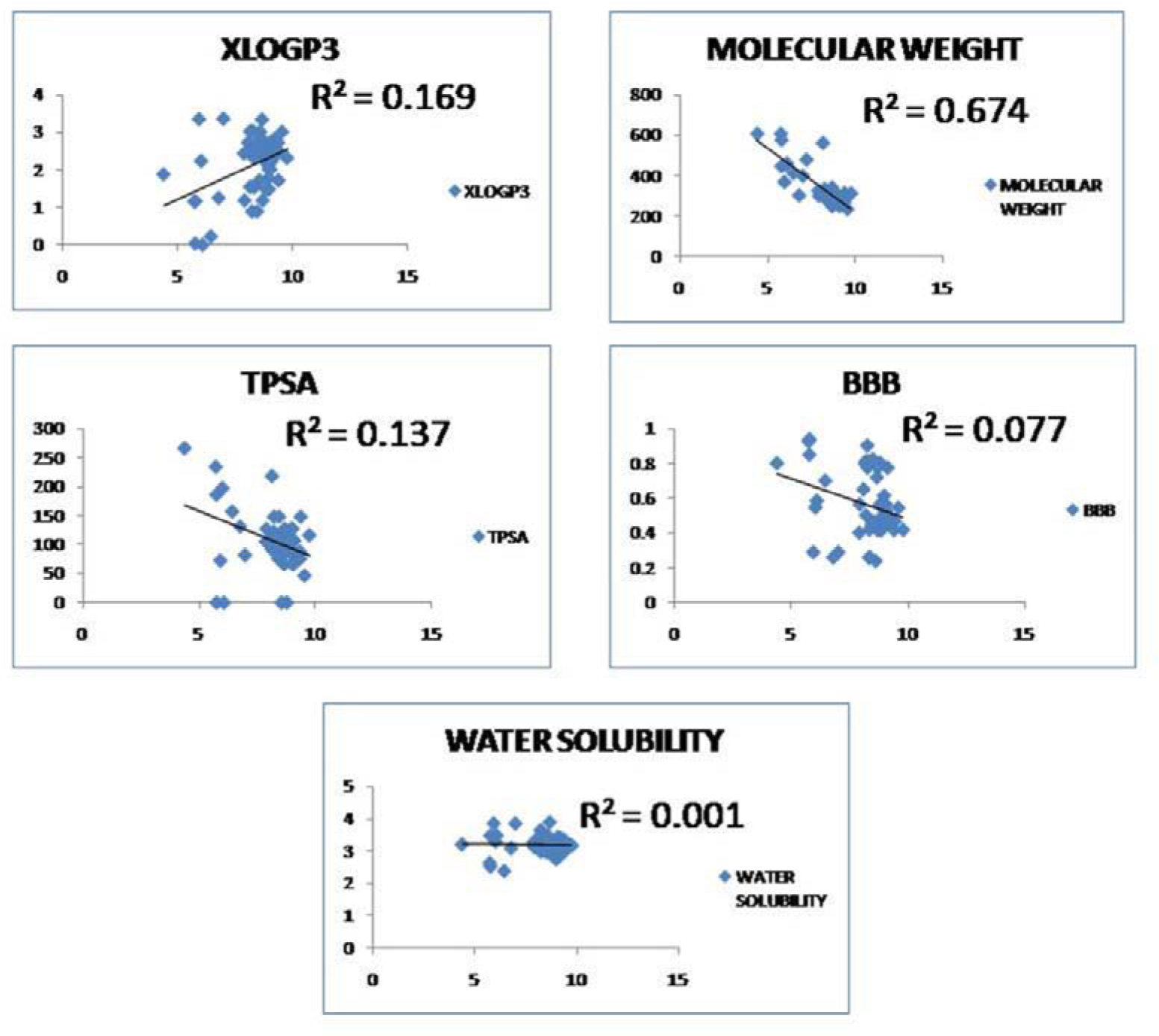
Plots of different pharmaco-kinetic properties of the flavonoids with Autodock Binding Free Energy Values.

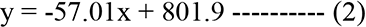

and

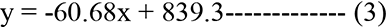

with the corresponding R² values, 0.690, 0.741 and 0.685 for (1), (2) and (3) respectively as can be seen in Figure 2 and Supplementary Table 3. These three equations were then used to predict the corresponding MW of the test set of flavonoids as grouped in the three sets (SET I, SET II and SET III). The observed MW and the predicted MW of the three sets were then plotted to check for the accuracy of the model as shown in Figure 2. The mean coefficient of determination R^2^ was calculated and was found to be 0.8 ± 0.11 from the three observations as can be seen in Figure 3. Cross-validation of the correlation obtained from the regression model was also performed for further precision. For internal validation, LOO cross-validation Q^2^ was found to be 0.674 (Supplementary Table 4). The external validation *R^2^* was found to be 0.7±0.045 (Supplementary Table 3).

**Figure 2:**
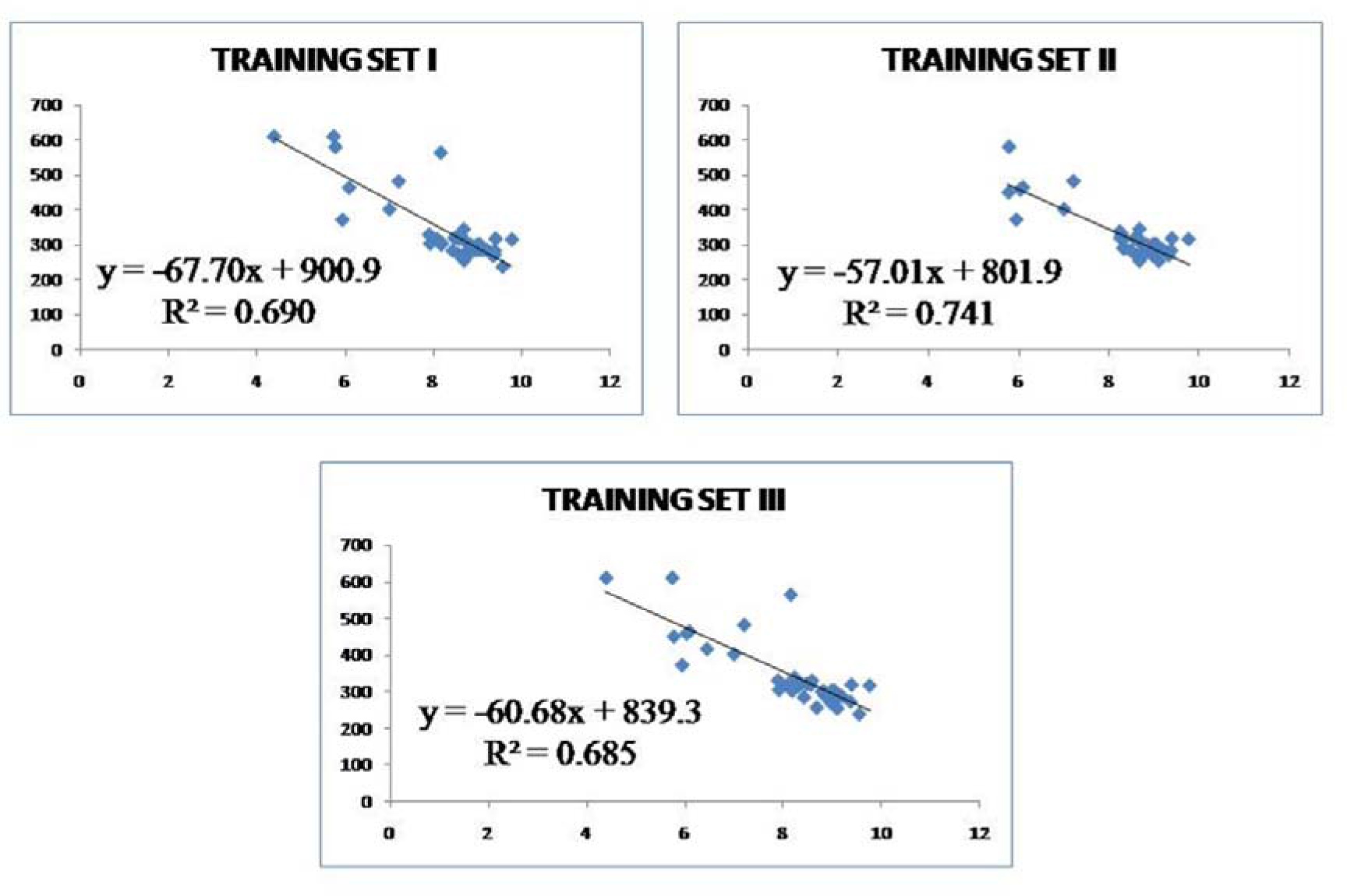
Three sets of Training group flavonoids relating Autodock binding free energy values with Molecular Weight of the flavonoids.

**Figure 3:**
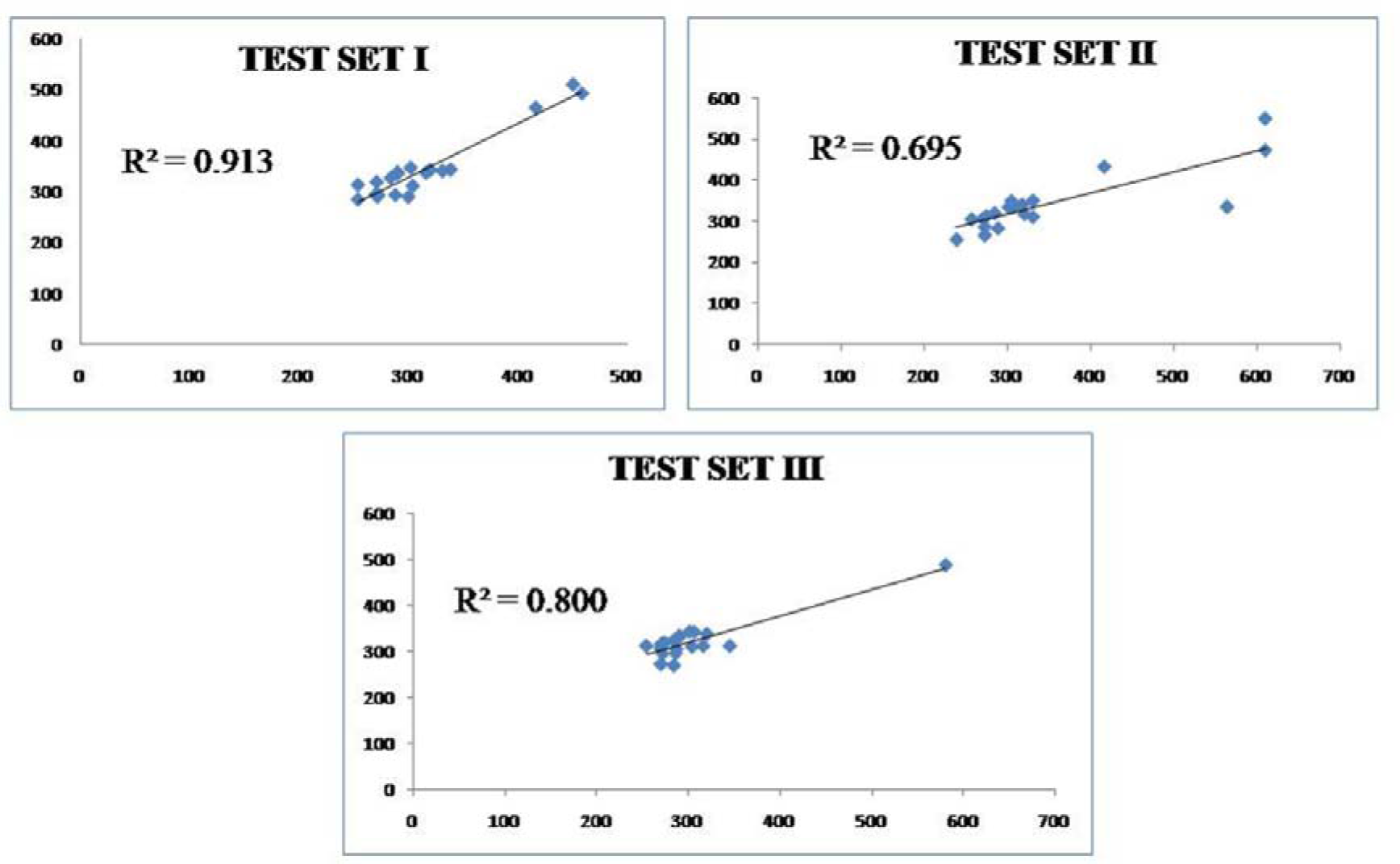
Graph showing observed Molecular weight vs. Predicted Molecular weight of the three sets of Test group flavonoids

In this section, we will discuss how the structure of the flavonoids would affect the binding free energies when they interact with the target Parkin protein. Hence, we first divided the flavonoids according to their backbone structure and compared how their structures differ from each other and how their binding free energy values change accordingly.

### Anthocyanin

Among anthocyanins, presence of different substituents at the position 5, 7, 3’, 4’, and 5’ brought about major changes in the binding free energy values with Parkin protein. The presence of –OH and –OCH_3_ groups at these positions affected their efficacies. The binding free energy was significant when –OH groups were present at positions 5, 7 and 4’ as in pelargonidin. Addition of another –OH group at position 3’ in case of cyanidin instead of – OCH_3_ group as in the case of peonidin, made the binding of the former more efficient, which showed the best binding among the anthocyanins. When six –OH groups were present at positions 5, 7, 3’, 4’, and 5’ as in delphinidin, the binding was good. When the –OH group at 5’ position of delphinidin was replaced with an –OCH_3_ group, the binding was lowered, as in petunidin. When another –OH group at position 3’ was replaced with the –OCH_3_ group, the structure became malvidin, where the binding free energy increased as compared to petunidin. However, when another –OCH_3_ group was added in place of –OH group at position 7, the structure became hirsutidin, the binding was significantly higher.

### Flavanone

In case of flavanones, the positions 5, 7, 3’, 4’ and 5’ were important. The binding was affected by the presence or absence of –OH and –OCH_3_ groups at these positions. The structures pinocembrin, naringenin, butin and eriodictyol were compared where there was an increase in the number of –OH groups. Pinocembrin had –OH groups at positions 5 and 7 in the A-ring, where the B ring had no substituent, yet the binding was significant. When an – OH group was added at position 4’ in the B ring, the binding was improved as in case of naringenin. However, when the three –OH groups were shuffled to positions 5, 4’ and 3’ in case of butin instead of 5,7 and 4’ in case of naringenin, there was a significant increase in the binding free energy of the flavanone with the protein. Butin showed the best binding to the Parkin protein among the flavanones. When another –OH group was added at position 3’ in case of eriodictyol as compared to naringenin, the binding was also significant. Addition of –OCH_3_ groups also showed some significant changes. The binding of sakuranetin to the protein was quite good when compared to naringenin, both of which differed by a –OCH_3_ group in their structures. However when both the –OCH_3_ and –OH groups were present in the B ring as in case of hesperetin instead of two –OH groups as in case of eriodictyol, (both of which had the same number of substients i.e. four), the binding was lowered. Addition of sugar residues at different positions as in cases of naringin, and hesperetin, decreased the binding as compared to their non-sugar counter parts, naringenin and hesperedin. Similar results were observed for astilibin where sugar residues at positions 3 decreased binding.

### Flavone

In this group, the best binder was 6-hydroxyflavone which was followed by wogonin, baicalein, 7,8 Dihydroxyflavone and chrysin. Notably, in all these structures presence of substituents in the A-ring and absence of substituents in the B-ring favored binding. Presence of hydroxyl groups at positions 5, 6, 7 and 8 in the A-ring and at positions 3’and 4’ in the B- ring favored binding. Presence of methoxy group at position 8 in A-ring was favored. However, increase of substituents in both A-ring and B-ring lowered the binding efficacy probably due to a crowding effect. 6-hydroxyflavone which had a single hydroxyl group at position 6 showed the best binding however the binding efficacy of luteolin, diosmetin, tricin, apigenin, which had substituents in both A-ring and B-ring, was lower. The binding was insignificant for nobiletin and tangeritin which might be due to the shielding effect as these structures had all their positions in the A-ring, i.e., 5,6,7,8 filled up.

### Flavonol

Among the flavonols, azaleatin was seen to be the best binder followed by gossypetin, quercetin and kaempferol. Structural analysis on the binding efficiency revealed that presence of hydroxyl groups at positions 3’, 4’, 5 and 7 favored binding. When kaempferol and kaempferide are compared, it was observed that the hydroxyl group at 4’ position increased binding. However, for rhamnetin and rhamnazin, the presence of –OCH_3_ group at position 3’ instead of –OH group showed an increase in binding. Similar inferences could be drawn when azaleatin was compared to quercetin, where –OCH_3_ group at position 5 (in case of azaleatin) instead of –OH group (in case of quercetin), increased the binding significantly. Presence of the –OCH_3_ group at position 3’ in case of isorhamnetin increased the binding than in case of rhamnetin in which it was at position 7. The binding efficacy of the flavonol glycosides used in docking i.e. myricitin and rutin, was insignificant where, the binding free energy of myricitin was higher than rutin.

### Isoflavonoid

The highest binding efficacy to Parkin protein, in case of isoflavonoids, was observed in case of daidzein followed by genistein, both of which had similar structures but genistein had an extra –OH group at position 5. However, the binding free energy decreased when the –OH group at position 4’ in genistein was replaced by a –OCH_3_ group in biochanin. This showed that –OH group at position 4’ favored binding. Addition of 4’ –OCH_3_ along with 5’ –OH groups in case of calycosin was found to have similar binding efficacy as biochanin. Notably, puerarin had a 4’ –OH group along with a glycosidic bond at the 8th position which lowered the binding free energy significantly.

### Chalcone

When we compared the chalcones like theaflavin, epigallocatechin, catechin, epicatechin and epigallocatechin-3-gallate (EGCG) we observed that the presence of –OH groups in the B ring can increase the binding efficacy. Epicatechin and catechin have similar structures; this leads to similar patterns of their interactions with Parkin as reflected by their similar binding free energy values. In contrast, EGCG and epigallocatechin, which differed only at position 3 of C-ring, their binding free energy values, are different. Epigallocatechin showed good binding efficiency than EGCG, which could be due to the branching in the C-ring of EGCG reduced binding. The best binder in this group was found to be the phloretin which had hydroxyl groups at positions 4, 2’, 4’ and 6’.

### Flavanonol

On comparing the structures of garbanzol and aromadedrin it was observed that, both the structures differed in only in one position, i.e., position 5 where there is an additional –OH group in aromadedrin. The binding efficacy of aromadedrin was comparatively lower than that of garbanzol. Garbanzol was the best binder among flavanonol group. When a –OH group was added to aromadedrin at position 3’, which converts it to taxifolin, the binding free energy value was lowered further. This lowering in the binding free energy values was consistent when another –OH group was added to position 2’ in case of dihydromorin and 3’ -OH group is shifted into the 4’ position. However, when five –OH groups were present in ampelopsin at positions 5, 7, 3’, 4’, and 5’ the binding efficacy was restored. When a –OH groups was added to the A ring (8^th^ position) as in taxifolin to dihydrogossypetin the binding free energy was decreased. Any branching in the rings reduces the binding free energy as in silibinin b.

## DISCUSSION

In order to create a trustworthy statistical model for the prediction of the biological activities of new chemical entities, QSAR have been used for decades to develop relationships between the physicochemical properties of chemical substances and their biological activities (Verma et al., 2010). Here, in this study we were interested in investigating the binding of 60 known bioactive flavonoids with the Parkin protein. In order to probe, whether this binding was related to their structural properties, structure-based QSAR was performed. Different pharmaco-kinetic properties of the flavonoids were calculated such as XLOGP3, MW, TPSA, BBB, water solubility and then these properties were compared with the Autodock B.F.E values of the 60 flavonoids to Parkin protein using regression analysis (Bellocchi et al., 2005) (Mitra et al., 2019). Figure 1 shows that among all the pharmaco-kinetic properties, MW of the flavonoids gave a co-efficient of determination value of 0.674. Generally, a R^2^ > 0.5 indicates a good correlation (Chicco et al., 2021). This correlation was further tested through training set-test set analysis. This experiment was repeated thrice to avoid biasness, each time with different combinations of training set and test set flavonoids as obtained from stratified random sampling. The three sets gave us three regression model equations with an average co-efficient of determination value of 0.705 ± 0.03 indicating a good linear relationship between the two parameters. A positive correlation between the MW of the flavonoids and Autodock B.F.E values of the flavonoids was obtained. The predicted MW of the test set flavonoids of the three sets (SET I, SET II and SET III) were then calculated from their corresponding regression model equations. The three corresponding sets of observed MW and the predicted MW of the test group flavonoids were then plotted (Figure 3). A mean coefficient of determination R^2^ was found to be 0.8 ± 0.11 from three observations which indicated good predictivity (Chicco et al., 2021). Cross-validation of the model was done both by internal and external cross-validation metrices to further test the predictivity of the model (Roy et al., 2015). For internal validation, Leave-one out mechanism was followed. The LOO cross-validation *Q^2^* was 0.674 (Supplementary Table 4). The external validation *R^2^* was found to be 0.7±0.045 (Supplementary Table 3). The internal validation metric LOO cross-validation *Q^2^* and the co-efficient of determination R^2^ values are highly significant. A model becomes overtrained, if the difference between these two parameters is more than 0.3 (Mitra et al., 2019). In our case it was found to be 0.126 which negates out such a possibility. The predictivity of the model is also assessed by the external validation metric *R^2^*_pred_ value. Generally, a good model follows the criteria - R^2^ > 0.8, *Q^2^* > 0.5, *R^2^* > 0.6 (Shamsara et al., 2017). Our model was found to satisfy all of these standard criteria thereby indicating its good predictivity. Therefore, we can clearly say that there was a good correlation between the MW of the flavonoids and their Autodock B.F.E. to the Parkin protein. Knowing any one of the parameters of the present or new group of flavonoids can help us predicting the other parameter with the help of our model.

## CONCLUSION

In summary, the presence of both the –OCH_3_ and -OH groups would help in making both polar and non-polar contacts with the amino acid residues of Parkin. Increase in the number of -OH groups would increase the extent of polar interactions. However, in such cases, the non-polar or hydrophobic interactions get diminished. At the same time, presence of more than one –OCH_3_ group in the flavonoids, though could increase the hydrophobic interactions, would enhance the steric clashes in the binding interface thereby destabilizing the binding interactions.

Our results could testify the hypothesis that flavonoid molecules can be used as suitable alternatives for the prevention and treatment of PD. The actual mechanism of action of flavonoids in PD is still not clear though. However, flavonoids modulate numerous important physiological responses, which eventually contribute to neuroprotection. Flavonoids may prevent and restore neuronal loss, and depletion of dopamine concentration. The flavonoids have their roles in improving the antioxidant status, reducing neuro-inflammation and mitochondrial dysfunction, inhibiting the α-synuclein aggregations to name of few. Among all the flavonoids, tested here, azaleatin shows the highest binding free energy of -9.77 Kcal/mol. The flavonoid has adequate distributions of hydrophobic and hydrophilic moieties. Therefore, it can easily fit into the active site cavity of Parkin as well as it can make polar contacts with the adjacent charged environment created by the polar amino acids present in the active site of Parkin.

Our work, for the first time, could come up with a molecular level view of the binding interactions of the flavonoids with Parkin. Such bindings of the flavonoids with Parkin would be able to protect the protein from acquiring mutations which result in the onset of PD. The main aim of the work is to come up with some alternative natural remedies to prevent the onset of PD by controlling the appearances of the causative mutations in the Parkin protein. Interestingly, most of the flavonoids, studied in this work, were found to be interacting with the amino acid residues which have already been found to be prone to acquire mutations leading to the onset of PD. This would further establish the protective actions of the flavonoids. Therefore, they may be used as suitable protective drug candidates against PD. This is the first report of the mechanistic details of the protective actions of flavonoids. These flavonoids may be tested further for their efficacies in the treatment of PD.

## AUTHORS’ CONTRIBUTION

SB: acquisition of data, performing the experiments, analysis and interpretation of data, drafting the article.

AM: performing the experiments, analysis and interpretation of data, drafting the article. SR: performing the experiments.

RG: revising the article critically for important intellectual content. AB: conceptualization of the work, drafting and revising the article. All the authors agreed to submit the manuscript in the current form.

## ACKNOWLEDGEMENT

The authors acknowledge the financial support from the DBT funded Bioinformatics Infrastructure Facility Center (Sanction no.: BT/PR40162/BTIS/137/48/2022 sanctioned to Prof. Angshuman Bagchi). Sima Biswas receives funding from University of Kalyani. Other funding sources are DST PURSE, SAP-UGC.

## REFERENCES

1. Adedayo, B. C., Oboh, G., Oyeleye, S. I., Ejakpovi, I. I., Boligon, A. A., & Athayde, M. L. (2015). Blanching alters the phenolic constituents and in vitro antioxidant and anticholinesterases properties of fireweed (Crassocephalum crepidioides). Journal of Taibah University Medical Sciences, 10(4), 419–426.

2. Alavijeh, M.S., Chishty, M., Qaiser, M. Z. & Palmer, A. M. (2005). Drug metabolism and Pharmacokinetics, the Blood-Brain Barrier, and Central Nervous System Drug Discovery. NeuroRx, 2(4), 554–571.

3. Alexander, G. E. (2004). Biology of Parkinson’s disease: pathogenesis and pathophysiology of a multisystem neurodegenerative disorder. Dialogues in clinical neuroscience, 6(3), 259.

4. Atrahimovich, D., Avni, D., & Khatib, S. (2021). Flavonoids-macromolecules interactions in human diseases with focus on Alzheimer, atherosclerosis and cancer. Antioxidants, 10(3), 423.

5. Ball, N., Teo, W. P., Chandra, S., & Chapman, J. (2019). Parkinson’s disease and the environment. Frontiers in Neurology, 10, 218.

6. Baron, J. A. (1996). Beneficial effects of nicotine and cigarette smoking: the real, the possible and the spurious. British Medical Bulletin, 52(1), 58–73.

7. Bellocchi, D., Macchiarulo, A., Costantino, G., & Pellicciari, R. (2005). Docking studies on PARP-1 inhibitors: insights into the role of a binding pocket water molecule. Bioorganic & medicinal chemistry, 13(4), 1151–1157.

8. Berman, H. M., Westbrook, J., Feng, Z., Gilliland, G., Bhat, T. N., Weissig, H., Shindyalov, I.N. & Bourne, P. E. (2000). The protein data bank. Nucleic acids research, 28(1), 235–242.

9. Blesa, J., Trigo-Damas, I., Quiroga-Varela, A., & Jackson-Lewis, V. R. (2015). Oxidative stress and Parkinson’s disease. Frontiers in Neuroanatomy, 9, 91.

10. Brooks, B. R., Brooks III, C. L., Mackerell Jr, A. D., Nilsson, L., Petrella, R. J., Roux, B., Won, Y., Archontis, G., Bartels, C., Boresch, S., Caves L, Cui Q, Dinner AR, Feig M, Fischer S, Gao J, Hodoscek M, Im W, Kuczera, K., Lazaridis, T., M, J., Ovchinnikov, V., Paci, E., Pastor, R. W., Post, C. B., Pu, J. Z., Schaefer, M., Tidor, B., Venable, R. M., Woodcock, H. L., Wu, X., Yang, W., York, D. M., Caflisch, A., & Karplus, M. (2009). CHARMM: the biomolecular simulation program. Journal of computational chemistry, 30(10), 1545–1614.

11. Cagac, A. (2020). Farming, well water consumption, rural living, and pesticide exposure in early life as the risk factors for Parkinson disease in Iğdır province. Neurosciences Journal, 25(2), 129–135.

12. Chai, C., & Lim, K. L. (2013). Genetic insights into sporadic Parkinson’s disease pathogenesis. Current Genomics, 14(8), 486–501.

13. Chicco, D., & Matthijs, J. (2021). Warrens, and Giuseppe Jurman.‘The coefficient of determination R-squared is more informative than SMAPE, MAE, MAPE, MSE and RMSE in regression analysis evaluation’. PeerJ Computer Science, 7, e623.

14. Chin-Chan, M., Navarro-Yepes, J., & Quintanilla-Vega, B. (2015). Environmental pollutants as risk factors for neurodegenerative disorders: Alzheimer and Parkinson diseases. Frontiers in Cellular Neuroscience, 9, 124.

15. Chung, K. K., Dawson, V. L., & Dawson, T. M. (2001). The role of the ubiquitin-proteasomal pathway in Parkinson’s disease and other neurodegenerative disorders. Trends in Neurosciences, 24, 7–14.

16. Collier, T. J., Kanaan, N. M., & Kordower, J. H. (2011). Ageing as a primary risk factor for Parkinson’s disease: evidence from studies of non-human primates. Nature Reviews Neuroscience, 12(6), 359–366.

17. Colovos, C., & Yeates, T. O. (1993). Verification of protein structures: patterns of nonbonded atomic interactions. Protein science, 2(9), 1511–1519.

18. Cookson, M. R. (2012). Parkinsonism due to mutations in PINK1, parkin, and DJ-1 and oxidative stress and mitochondrial pathways. Cold Spring Harbor Perspectives in Medicine, 2(9), a009415.

19. de Andrade Teles, R. B., Diniz, T. C., Costa Pinto, T. C., de Oliveira Junior, R. G., Gama e Silva, M., de Lavor, É. M., Fernandes, A.W.C., de Oliveira, A.P., de Almeida Ribeiro, F.P.R., da Silva, A.A.M., Cavalcante, T.C.F., Quintans Júnior, L.J., & da Silva Almeida, J. R. G. (2018). Flavonoids as therapeutic agents in Alzheimer’s and Parkinson’s diseases: a systematic review of preclinical evidences. Oxidative medicine and cellular longevity, 2018.

20. DeMaagd, G., & Philip, A. (2015). Parkinson’s disease and its management: part 1: disease entity, risk factors, pathophysiology, clinical presentation, and diagnosis. Pharmacy and therapeutics, 40(8), 504.

21. Eisenberg, D., Lüthy, R., & Bowie, J. U. (1997). [20] VERIFY3D: assessment of protein models with three-dimensional profiles. In Methods in enzymology (Vol. 277, pp. 396–404). Academic Press.

22. Emamzadeh, F. N., & Surguchov, A. (2018). Parkinson’s disease: biomarkers, treatment, and risk factors. Frontiers in neuroscience, 612.

23. Falcone Ferreyra, M. L., Rius, S., & Casati, P. (2012). Flavonoids: biosynthesis, biological functions, and biotechnological applications. Frontiers in plant science, 3, 222.

24. Gabler, F., Nam, S. Z., Till, S., Mirdita, M., Steinegger, M., Söding, J., Lupas, A.N. & Alva, V. (2020). Protein sequence analysis using the MPI bioinformatics toolkit. Current Protocols in Bioinformatics, 72(1), e108.

25. Gasteiger, E., Hoogland, C., Gattiker, A., Wilkins, M. R., Appel, R. D., & Bairoch, A. (2005). Protein identification and analysis tools on the ExPASy server. The proteomics protocols handbook, 571-607.

26. Gatto, N. M., Cockburn, M., Bronstein, J., Manthripragada, A. D., & Ritz, B. (2009). Well-water consumption and Parkinson’s disease in rural California. Environmental Health Perspectives, 117(12), 1912–1918.

27. Gómez-Benito, M., Granado, N., García-Sanz, P., Michel, A., Dumoulin, M., & Moratalla, R. (2020). Modeling Parkinson’s disease with the alpha-synuclein protein. Frontiers in Pharmacology, 11, 356.

28. Guo, J. F., Xiao, B., Liao, B., Zhang, X. W., Nie, L. L., Zhang, Y. H., Shen, L., Jiang, H., Xia, K., Pan, Q., Yan, X.X & Tang, B. S. (2008). Mutation analysis of Parkin, PINK1, DJ-1 and ATP13A2 genes in Chinese patients with autosomal recessive early-onset Parkinsonism. Movement disorders: official journal of the Movement Disorder Society, 23(14), 2074–2079.

29. Halgren, T. A. (1996). Merck molecular force field. I. Basis, form, scope, parameterization, and performance of MMFF94. Journal of computational chemistry, 17(5-6), 490–519.

30. Hernán, M. A., Takkouche, B., Caamaño-Isorna, F., & Gestal-Otero, J. J. (2002). A meta-analysis of coffee drinking, cigarette smoking, and the risk of Parkinson’s disease. Annals of Neurology, 52(3), 276–284.

31. Hindle, J. V. (2010). Ageing, neurodegeneration and Parkinson’s disease. Age and Ageing, 39(2), 156–161.

32. Hussein, R. A., & El-Anssary, A. A. (2019). Plants secondary metabolites: the key drivers of the pharmacological actions of medicinal plants. Herbal Medicine, 1, 13.

33. Jung, U. J., & Kim, S. R. (2018). Beneficial effects of flavonoids against Parkinson’s disease. Journal of medicinal food, 21(5), 421–432.

34. Kouli, A., Torsney, K. M., & Kuan, W. L. (2018). Parkinson’s disease: etiology, neuropathology, and pathogenesis. Exon Publications, 3-26.

35. Kumar, S., & Pandey, A. K. (2013). Chemistry and biological activities of flavonoids: an overview. The scientific world journal, 2013.

36. Li, J. Q., Tan, L., & Yu, J. T. (2014). The role of the LRRK2 gene in Parkinsonism. Molecular Neurodegeneration, 9(1), 1–17.

37. Lipinski, C. A. (2004). Lead-and drug-like compounds: the rule-of-five revolution. Drug discovery today: Technologies, 1(4), 337–341.

38. Magalingam, K. B., Radhakrishnan, A. K., & Haleagrahara, N. (2015). Protective mechanisms of flavonoids in Parkinson’s disease. Oxidative medicine and cellular longevity, 2015.

39. Mitra, A., Biswas, R., Bagchi, A., & Ghosh, R. (2019). Insight into the binding of a synthetic nitro-flavone derivative with human poly (ADP-ribose) polymerase 1. International journal of biological macromolecules, 141, 444–459.

40. Nuytemans, K., Theuns, J., Cruts, M., & Van Broeckhoven, C. (2010). Genetic etiology of Parkinson disease associated with mutations in the SNCA, PARK2, PINK1, PARK7, and LRRK2 genes: a mutation update. Human Mutation, 31(7), 763-780.

41. Panche, A. N., Diwan, A. D., & Chandra, S. R. (2016). Flavonoids: an overview. Journal of Nutritional Science, 5.

42. Pang, S. Y. Y., Ho, P. W. L., Liu, H. F., Leung, C. T., Li, L., Chang, E. E. S., Ramsden, D.B. & Ho, S. L. (2019). The interplay of aging, genetics and environmental factors in the pathogenesis of Parkinson’s disease. Translational Neurodegeneration, 8(1), 1–11.

43. Pardridge, W. M. (2012). Drug transport across the blood–brain barrier. Journal of cerebral blood flow & metabolism, 32(11), 1959–1972.

44. Parkinson, J. (2002). An essay on the shaking palsy. The Journal of neuropsychiatry and clinical neurosciences, 14(2), 223–236.

45. Picca, A., Calvani, R., Coelho-Junior, H. J., Landi, F., Bernabei, R., & Marzetti, E. (2020). Mitochondrial dysfunction, oxidative stress, and neuroinflammation: Intertwined roads to neurodegeneration. Antioxidants, 9(8), 647.

46. Ramachandran, G. T., & Sasisekharan, V. (1968). Conformation of polypeptides and proteins. Advances in protein chemistry, 23, 283–437.

47. Ren, X., & Chen, J. F. (2020). Caffeine and Parkinson’s disease: multiple benefits and emerging mechanisms. Frontiers in Neuroscience, 1334.

48. Roy, K., Kar, S., Das, R. N., Roy, K., Kar, S., & Das, R. N. (2015). Statistical methods in QSAR/QSPR. A Primer on QSAR/QSPR Modeling: Fundamental Concepts, 37-59.

49. Shamsara, J. (2017). Ezqsar: an R package for developing QSAR models directly from structures. The Open Medicinal Chemistry Journal, 11, 212.

50. Song, J., & Kim, J. (2016). Degeneration of dopaminergic neurons due to metabolic alterations and Parkinson’s disease. Frontiers in Aging Neuroscience, 8, 65.

51. Spataro, N., Calafell, F., Cervera-Carles, L., Casals, F., Pagonabarraga, J., Pascual-Sedano, B., Campolongo, A., Kulisevsky, J., Lleó, A., Navarro, A., Clarimón, J., & Bosch, E. (2015). Mendelian genes for Parkinson’s disease contribute to the sporadic forms of the disease. Human Molecular Genetics, 24(7), 2023-2034.

53. Stefanis, L. (2012). α-Synuclein in Parkinson’s disease. Cold Spring Harbor perspectives in medicine, 2(2), a009399.

54. Tan, E. K., Tan, C., Fook-Chong, S. M. C., Lum, S. Y., Chai, A., Chung, H., Shen, H., Zhao, Y., Teoh, M.L., Yih, Y., Pavanni, R., Chandran, V. R. & Wong, M. C. (2003). Dose-dependent protective effect of coffee, tea, and smoking in Parkinson’s disease: a study in ethnic Chinese. Journal of the Neurological Sciences, 216(1), 163–167.

55. Thilakarathna, S. H., & Rupasinghe, H. P. (2013). Flavonoid bioavailability and attempts for bioavailability enhancement. Nutrients, 5(9), 3367–3387.

56. Triarhou, L. C. (2013). Dopamine and Parkinson’s disease. In Madame Curie Bioscience Database [Internet]. Landes Bioscience.

57. Ullah, A., Munir, S., Badshah, S. L., Khan, N., Ghani, L., Poulson, B. G., Emwas, A.H. & Jaremko, M. (2020). Important flavonoids and their role as a therapeutic agent. Molecules, 25(22), 5243.

58. Verma, J. “KhedkarVM and Coutinho EC. 3D-QSAR in drug design--a review.” Current Topics in Medicinal Chemistry 10 (2010): 95–115.

59. Webb, B., & Sali, A. (2016). Comparative protein structure modeling using MODELLER. Current protocols in bioinformatics, 54(1), 5–6.

60. Yang, H., Lou, C., Sun, L., Li, J., Cai, Y., Wang, Z., Li, W., Liu, G. & Tang, Y. (2019). admetSAR 2.0: web-service for prediction and optimization of chemical ADMET properties. Bioinformatics, 35(6), 1067–1069.

61. Yang, X., & Xu, Y. (2014). Mutations in the ATP13A2 gene and Parkinsonism: a preliminary review. BioMed Research International, 2014.

62. Yonekura-Sakakibara, K., Higashi, Y., & Nakabayashi, R. (2019). The origin and evolution of plant flavonoid metabolism. Frontiers in Plant Science, 10, 943.

63. Zahoor, I., Shafi, A., & Haq, E. (2018). Pharmacological treatment of Parkinson’s disease. Exon Publications, 129–144.

64. Zimmermann, L., Stephens, A., Nam, S. Z., Rau, D., Kübler, J., Lozajic, M., Gabler, F., Söding, J., Lupas, A.N. & Alva, V. (2018). A completely reimplemented MPI bioinformatics toolkit with a new HHpred server at its core. Journal of molecular biology, 430(15), 2237–2243.

