## Supplementary figures and tables for "Molecular insight into the mechanism of action of some beneficial flavonoids for the treatment of Parkinson’s Disease"

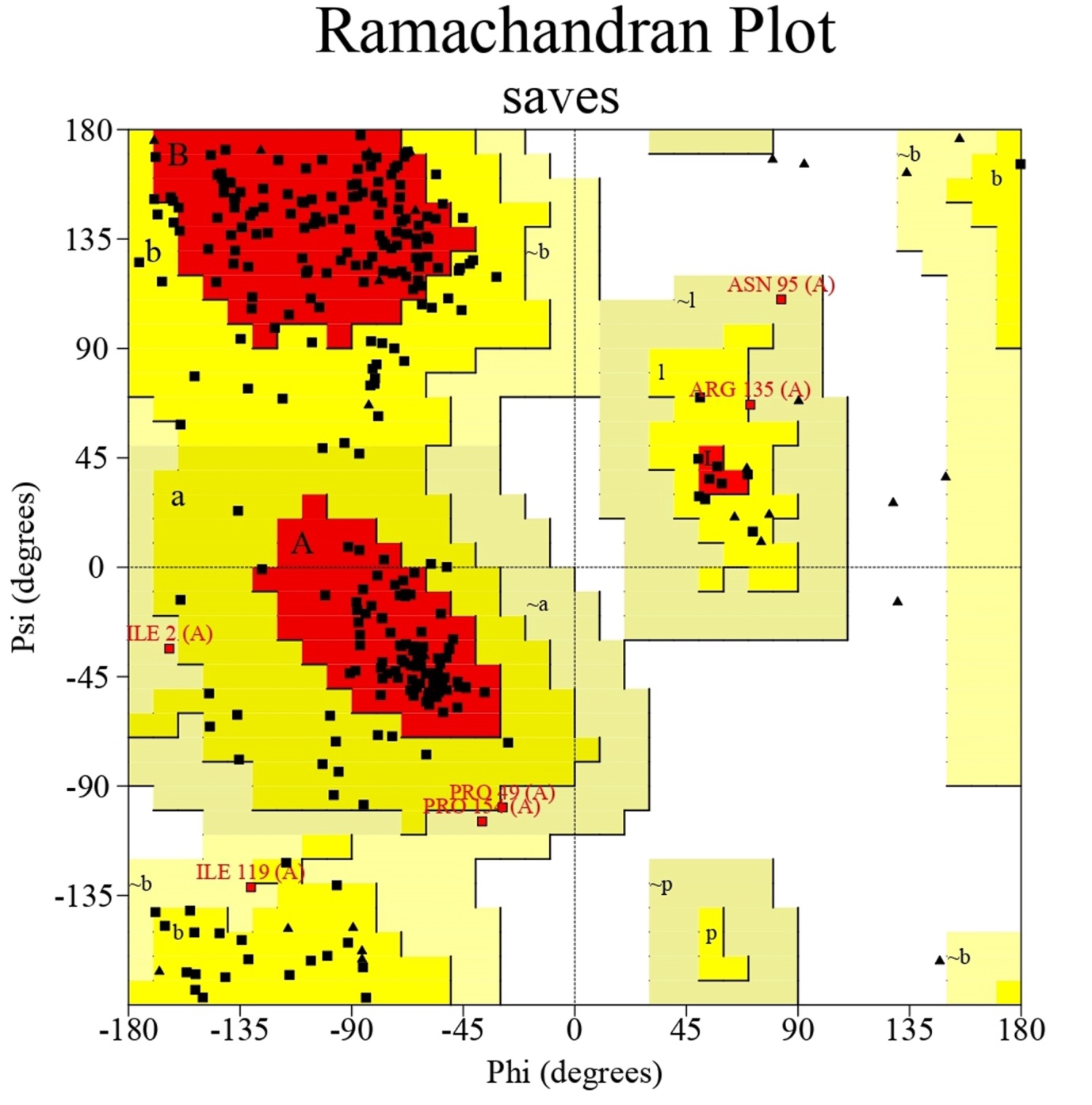

Supplementary Figure1: Ramachandran plot analysis of Parkin. In this plot, the most favoured region is marked in red; additionally allowed regions are marked in yellow; disallowed regions are white. No amino acid residues of Parkin were found to be present in the disallowed region of the Ramachandran plot.

Supplementary Table1: The details of the Lipiniski’s filter analyses of the selected flavonoids. Tables1.1-1.7: The results are presented as per the structural groupings of the selected flavonoids.

Supplementary Table 1.1:

|  | **Pelargonidin** | **Peonidin** | **Cyanidin** | **Delphinidin** | **Hirsutidin** | **Malvidin** | **Petunidin** |
| --- | --- | --- | --- | --- | --- | --- | --- |
| **Molecular weight** | **271** | **301** | **287** | **303** | **345** | **331** | **317** |
| **Num. H-bond acceptors** | **5** | **6** | **6** | **7** | **7** | **7** | **7** |
| **Num. H-bond donors** | **4** | **4** | **5** | **6** | **3** | **4** | **5** |
| **LOGP** | **3.013289** | **3.021889** | **2.718889** | **2.42449** | **3.333489** | **3.03049** | **2.72749** |
| **Molar Refractivity** | **72.055176** | **78.607178** | **73.719978** | **75.384773** | **90.046371** | **85.159172** | **80.271973** |

Supplementary Table 1.2:

|  | **Eriodictyol** | **Butin** | **Sakuranetin** | **Naringenin** | **Naringin** | **Hesperetin** | **Hesperedin** | **Asitibilin** | **Pinocembrin** |
| --- | --- | --- | --- | --- | --- | --- | --- | --- | --- |
| **Molecular weight** | **288** | **272** | **286** | **272** | **580** | **302** | **610** | **450** | **256** |
| **Num. H-bond acceptors** | **6** | **5** | **5** | **5** | **14** | **6** | **15** | **11** | **4** |
| **Num. H-bond donors** | **4** | **3** | **2** | **3** | **8** | **3** | **8** | **7** | **2** |
| **LOGP** | **2.215499** | **2.509899** | **2.812899** | **2.509899** | **-1.1652** | **2.518499** | **-1.1566** | **0.038101** | **2.804299** |
| **Molar Refractivity** | **71.85968** | **70.194885** | **75.082077** | **70.194878** | **134.146881** | **76.74688** | **140.698914** | **104.471046** | **68.530083** |

Supplementary Table 1.3:

|  | **Apigenin** | **Baicalein** | **6-hydroxyflavone** | **Luteolin** | **Chrysin** | **Wogonin** | **Nobiletin** | **Diosmetin** | **Tricin** | **Tangeretin** | **7,8-dihydroxyflavone** |
| --- | --- | --- | --- | --- | --- | --- | --- | --- | --- | --- | --- |
| **Molecular weight** | **270** | **270** | **238** | **286** | **254** | **284** | **402** | **300** | **330** | **372** | **254** |
| **Num. H-bond acceptors** | **5** | **5** | **3** | **6** | **4** | **5** | **8** | **6** | **7** | **7** | **4** |
| **Num. H-bond donors** | **3** | **3** | **1** | **4** | **2** | **2** | **0** | **3** | **3** | **0** | **2** |
| **LOGP** | **2.419599** | **2.419599** | **3.008399** | **2.125199** | **2.713999** | **2.722598** | **3.354399** | **2.428199** | **2.4368** | **3.345799** | **2.713999** |
| **Molar Refractivity** | **70.813881** | **70.813881** | **67.484283** | **72.478676** | **69.149078** | **75.701073** | **105.131462** | **77.365875** | **83.91787** | **98.579468** | **69.149078** |

Supplementary Table 1.4:

|  | **Isorhamnetin** | **Fustin** | **Rutin** | **Kaempferol** | **Myricetin** | **Myricitrin** | **Azaleatin** | **Quercetin** | **Fisetin** | **Morin** | **Galangin** | **Gossypetin** | **Kaempferide** | **Rhamnetin** | **Rhamnazin** |
| --- | --- | --- | --- | --- | --- | --- | --- | --- | --- | --- | --- | --- | --- | --- | --- |
| **Molecular weight** | **316** | **288** | **610** | **286** | **318** | **464** | **316** | **302** | **286** | **302** | **270** | **318** | **300** | **316** | **330** |
| **Num. H-bond acceptors** | **7** | **6** | **16** | **6** | **8** | **12** | **7** | **7** | **6** | **7** | **5** | **8** | **6** | **7** | **7** |
| **Num. H-bond donors** | **4** | **4** | **10** | **4** | **6** | **8** | **4** | **5** | **4** | **5** | **3** | **6** | **3** | **4** | **3** |
| **LOGP** | **2.3139** | **1.4807** | **-1.8788** | **2.305299** | **1.7165** | **0.0026** | **2.313899** | **2.0109** | **2.3053** | **2.0109** | **2.599699** | **1.7165** | **2.608299** | **2.313899** | **2.616899** |
| **Molar Refractivity** | **78.937675** | **71.584679** | **137.495483** | **72.385681** | **75.715279** | **106.52684** | **78.937675** | **74.050476** | **72.385681** | **74.050476** | **70.720879** | **75.715271** | **77.272873** | **78.937675** | **83.824867** |

Supplementary Table 1.5:

|  | **Daidzein** | **Biochanin** | **Genistein** | **Calycosin** | **Puerarin** |
| --- | --- | --- | --- | --- | --- |
| **Molecular weight** | **254** | **284** | **270** | **248** | **416** |
| **Num. H-bond acceptors** | **4** | **5** | **5** | **5** | **9** |
| **Num. H-bond donors** | **2** | **2** | **3** | **2** | **6** |
| **LOGP** | **2.713999** | **2.722599** | **2.419599** | **2.722599** | **0.2289** |
| **Molar Refractivity** | **69.149086** | **75.70108** | **70.813881** | **75.70108** | **101.869247** |

Supplementary Table 1.6:

|  | **Catechin** | **Epicatechin** | **Epigallocatechin** | **Theaflavin** | **EGCG** | **Phloretin** |
| --- | --- | --- | --- | --- | --- | --- |
| **Molecular weight** | **290** | **290** | **306** | **566** | **458** | **274** |
| **Num. H-bond acceptors** | **6** | **6** | **7** | **12** | **11** | **5** |
| **Num. H-bond donors** | **5** | **5** | **6** | **9** | **8** | **4** |
| **LOGP** | **1.5461** | **1.5461** | **1.2517** | **1.348** | **2.233201** | **2.324499** |
| **Molar Refractivity** | **72.622978** | **72.622978** | **74.287773** | **138.645706** | **108.920845** | **72.194679** |

Supplementary Table 1.7:

| **Molecular weight** | **Taxifolin** | **Dihydrogossypetin** | **Dihydromorin** | **Garbanzol** | **Aromadedrin** | **Ampelopsin** | **Silibinin B** |
| --- | --- | --- | --- | --- | --- | --- | --- |
| **Num. H-bond acceptors** | **304** | **320** | **304** | **272** | **288** | **320** | **482** |
| **Num. H-bond donors** | **7** | **8** | **7** | **5** | **6** | **8** | **10** |
| **LOGP** | **5** | **6** | **5** | **3** | **4** | **6** | **5** |
| **Molar Refractivity** | **1.1863** | **0.8919** | **1.1863** | **1.7751** | **1.4807** | **0.8919** | **2.3627** |
|  | **73.249474** | **74.914268** | **73.249474** | **69.919884** | **71.584679** | **74.914276** | **119.44944** |

Supplementary Table2: The details of the ADMET profiles of the selected flavonoids. The data are presented in the excel spreadsheet marked as Drug_likeliness_properties_ADMETSAR.

Supplementary Table 3: Three sets of training and test groups of flavonoids and the corresponding calculated values of parameters for external cross-validation of the correlation relating Autodock binding free energy with Molecular Weight of the flavonoids obtained from regression analysis.

| **TRAINING SET OF FLAVONOIDS** | **TEST SET OF FLAVONOIDS** | **MODEL EQUATION OBTAINED** | | **R^2^** | | **PREDICTING FOR THE TEST SET OF FLAVONOIDS** | | | | |
| --- | --- | --- | --- | --- | --- | --- | --- | --- | --- | --- |
|  |  |  |  |  |  | ***{Y_obs(test)-_ Y_pred(test)_}^2^*** | ***{Y_obs(test)-_ Ȳ_(train)_ }^2^*** | | **R^2^ _pred_** | |
| **SET I** | | | | | | | | | | |
| **1** \| **PETUNIDIN**; **2** \| **PEONIDIN**; **3** \| **CYANIDIN**; **4** \| **HIRSUTIDIN**; **5** \| **THEAFLAVIN**; **6** \| **PHLORETIN**; **7** \| **EIPGALLOCATECHIN**; **8** \| **NARINGIN**; **9** \| **ERIODICTYOL**; **10** \| **PINOCEMBRIN**; **11** \| **HESPERIDIN**; **12** \| **BUTIN**; **13** \| **SAKURANETIN**; **14** \| **AROMADEDRIN**; **15** \| **DIHYDROMORIN**; **16** \| **DIHYDROGOSSYPETIN**; **17** \| **TRICIN**; **18** \| **6-HYDROXYFLAVONE**; **19** \| **TANGERITIN**; **20** \| **BAICALEIN**; **21** \| **LUTEOLIN**; **22** \| **WOGONIN**; **23** \| **NOBILETIN**; **24** \| **DIOSMETIN**; **25** \| **MYRICITRIN**; **26** \| **ISORHAMNETIN**; **27** \| **GALANGIN**; **28** \| **MYRICETIN**; **29** \| **KAEMPFERIDE**; **30** \| **QUERCETIN**; **31** \| **KAEMPFEROL**; **32** \| **GOSSYPETIN**; **33** \| **MORIN**; **34** \| **AZALEATIN**; **35** \| **FISETIN**; **36** \| **RHAMNAZIN**; **37** \| **RUTIN**; **38** \| **SILIBININ B**; **39** \| **BIOCHANIN**; **40** \| **GENISTEIN** | **PELARGONIDIN**; **MALVIDIN**; **DELPHINIDIN**; **CATECHIN**; **EPICATECHIN**; **EGCG**; **NARINGENIN**; **HESPERETIN**; **ASTILBIN**; **TAXIFOLIN**; **AMPELOPSIN**; **GARBANZOL**; **7,8- DIHYDROXYFLAVONE**; **CHRYSIN**; **APIGENIN**; **FUSTIN**; **RHAMNETIN**; **DAIDZEIN**; **PUERARIN**; **CALYCOSIN** | **y = -67.70x + 900.9** | | **0.690** | | **2250.5536**; **81.793936**; **19.027044**; **2243.537956**; **2055.262225**; **1174.364361**; **323.856016**; **1949.840649**; **3584.536641**; **48.762289**; **489.515625**; **486.555364**; **894.787569**; **114.297481**; **547.138881**; **22.127616**; **428.448601**; **3483.832576**; **2290.101025**; **1809.141156** | **4543.541874**; **53.96077764**; **0.00195364**; **2340.218026**; **2340.218026**; **14341.06842**; **4408.402258**; **1322.47141**; **12489.00122**; **1183.071058**; **338.4054576**; **4408.402258**; **7124.339074**; **1473.469642**; **4679.353474**; **2539.736658**; **501.1240416**; **7124.339074**; **6042.60585**; **2956.727626** | | **0.697078** | |
| **SET II** | | | | | | | | | | |
| **1** \| **CYANIDIN**; **2** \| **HIRSUTIDIN**; **3** \| **PELARGONIDIN**; **4** \| **MALVIDIN**; **5** \| **DELPHINIDIN**; **6** \| **CATECHIN**; **7** \| **EPICATECHIN**; **8** \| **EGCG**; **9** \| **ASTILBIN**; **10** \| **NARINGIN**; **11** \| **SAKURANETIN**; **12** \| **TAXIFOLIN**; **13** \| **AMPELOPSIN**; **14** \| **GARBANZOL**; **15** \| **AROMADEDRIN**; **16** \| **TANGERITIN**; **17** \| **BAICALEIN**; **18** \| **LUTEOLIN**; **19** \| **WOGONIN**; **20** \| **NOBILETIN**; **21** \| **7,8- DIHYDROXYFLAVONE**; **22** \| **CHRYSIN**; **23** \| **APIGENIN**; **24** \| **DIOSMETIN**; **25** \| **FUSTIN**; **26** \| **RHAMNETIN**; **27** \| **MYRICITRIN**; **28** \| **ISORHAMNETIN**; **29** \| **GALANGIN**; **30** \| **MYRICETIN**; **31** \| **KAEMPFERIDE**; **32** \| **QUERCETIN**; **33** \| **KAEMPFEROL**; **34** \| **GOSSYPETIN**; **35** \| **MORIN**; **36** \| **AZALEATIN**; **37** \| **FISETIN**; **38** \| **SILIBININ B**; **39** \| **DAIDZEIN**; **40** \| **CALYCOSIN** | **PETUNIDIN**; **PEONIDIN**; **THEAFLAVIN**; **PHLORETIN**; **EIPGALLOCATECHIN**; **NARINGENIN**; **HESPERETIN**; **ERIODICTYOL**; **PINOCEMBRIN**; **HESPERIDIN**; **BUTIN**; **DIHYDROMORIN**; **DIHYDROGOSSYPETIN**; **TRICIN**; **6-HYDROXYFLAVONE**; **RHAMNAZIN**; **RUTIN**; **BIOCHANIN**; **GENISTEIN**; **PUERARIN** | **y = -57.01x + 801.9** | | **0.741** | | **575.4817166**; **1215.101079**; **51893.56896**; **1570.885663**; **857.7986592**; **237.770232**; **1069.819806**; **16.00640064**; **2522.359774**; **18470.82137**; **26.04877444**; **2128.050709**; **1.49989009**; **450.755361**; **347.6136514**; **327.8236148**; **3468.491457**; **1372.383888**; **1484.060052**; **317.0358303** | **45.857275**; **518.55488**; **57820.146**; **2478.2276**; **315.83688**; **2682.3905**; **473.57594**; **1281.0529**; **4594.3724**; **82098.409**; **2682.3905**; **391.71535**; **14.377747**; **39.040003**; **7361.9489**; **39.040003**; **82069.759**; **1582.5916**; **2894.6337**; **8526.3432** | | **0.657426** | |
| **SET III** | | | | | | | | | | |
| **1** \| **PETUNIDIN**; **2** \| **MALVIDIN**; **3** \| **DELPHINIDIN**; **4** \| **THEAFLAVIN**; **5** \| **EGCG**; **6** \| **NARINGENIN**; **7** \| **HESPERETIN**; **8** \| **ASTILBIN**; **9** \| **ERIODICTYOL**; **10** \| **PINOCEMBRIN**; **11** \| **HESPERIDIN**; **12** \| **BUTIN**; **13** \| **SAKURANETIN**; **14** \| **AROMADEDRIN**; **15** \| **DIHYDROMORIN**; **16** \| **DIHYDROGOSSYPETIN**; **17** \| **TRICIN**; **18** \| **6-HYDROXYFLAVONE**; **19** \| **TANGERITIN**; **20** \| **NOBILETIN**; **21** \| **7,8- DIHYDROXYFLAVONE**; **22** \| **CHRYSIN**; **23** \| **APIGENIN**; **24** \| **DIOSMETIN**; **25** \| **FUSTIN**; **26** \| **RHAMNETIN**; **27** \| **MYRICITRIN**; **28** \| **MYRICETIN**; **29** \| **KAEMPFERIDE**; **30** \| **QUERCETIN**; **31** \| **KAEMPFEROL**; **32** \| **GOSSYPETIN**; **33** \| **MORIN**; **34** \| **AZALEATIN**; **35** \| **FISETIN**; **36** \| **RHAMNAZIN**; **37** \| **RUTIN**; **38** \| **SILIBININ B**; **39** \| **BIOCHANIN**; **40** \| **PUERARIN** | **PEONIDIN**; **CYANIDIN**; **HIRSUTIDIN**; **PELARGONIDIN**; **PHLORETIN**; **EIPGALLOCATECHIN**; **CATECHIN**; **EPICATECHIN**; **NARINGIN**; **TAXIFOLIN**; **AMPELOPSIN**; **GARBANZOL**; **BAICALEIN**; **LUTEOLIN**; **WOGONIN**; **ISORHAMNETIN**; **GALANGIN**; **GENISTEIN**; **DAIDZEIN**; **CALYCOSIN** | | **y = -60.68x + 839.3** | | **0.685** | **1787.125**; **349.047**; **1070.755**; **2135.549**; **2081.111**; **1344.513**; **1951.201**; **1793.692**; **8451.198**; **42.60434**; **340.0336**; **545.5588**; **8.500723**; **99.48068**; **199.8944**; **13.41317**; **1361.344**; **1951.696**; **3405.609**; **1637.303** | | **1809.737**; **3200.278**; **2.277081**; **5266.55**; **4837.342**; **1409.327**; **2866.639**; **2866.639**; **56021.68**; **1565.073**; **555.1207**; **5120.977**; **5412.692**; **3314.42**; **3546.322**; **759.0576**; **5412.692**; **5412.692**; **8022.964**; **3545.131** | | **0.747249** |

Supplementary Table 3: Calculated values of parameters for internal cross-validation by Leave-One-Out mechanism of the correlation relating Autodock binding free energy with Molecular Weight of the flavonoids obtained from regression analysis.

| Flavonoids | Yobs(train) | Ŷtrain | Ypred(train) from LOO | {Yobs(train) - Ypred(train)}^2^ | {Yobs(train) - Ŷtrain}^2^ | PRESS | Q^2^ |
| --- | --- | --- | --- | --- | --- | --- | --- |
| CYANIDIN | **287.24** | **330.5303** | **297.7016** | **109.4451** | **1874.05** | **147106.3** | **0.674** |
| HIRSUTIDIN | **345.32** |  | **303.83** | **1721.42** | **218.7352** |  |  |
| PELARGONIDIN | **271.24** |  | **310.604** | **1549.524** | **3515.34** |  |  |
| MALVIDIN | **331.3** |  | **330.6516** | **0.420423** | **0.592438** |  |  |
| DELPHINIDIN | **338.69** |  | **333.1528** | **30.66058** | **66.5807** |  |  |
| PEONIDIN | **301.27** |  | **338.4147** | **1379.729** | **856.1652** |  |  |
| PETUNIDIN | **317.27** |  | **344.192** | **724.7941** | **175.8356** |  |  |
| BUTIN | **272.25** |  | **258.6426** | **185.1613** | **3396.593** |  |  |
| SAKURANETIN | **286.28** |  | **269.7317** | **273.8462** | **1958.089** |  |  |
| ERIODICTYOL | **288.25** |  | **278.4196** | **96.63676** | **1787.624** |  |  |
| NARINGENIN | **272.25** |  | **282.7314** | **109.8597** | **3396.593** |  |  |
| PINOCEMBRIN | **256.26** |  | **304.8487** | **2360.862** | **5516.077** |  |  |
| HESPERETIN | **302.28** |  | **337.1767** | **1217.78** | **798.0795** |  |  |
| NARINGIN | **580.5** |  | **483.496** | **9409.776** | **62484.85** |  |  |
| ASTILBIN | **450.4** |  | **498.8295** | **2345.416** | **14368.74** |  |  |
| HESPERIDIN | **610.57** |  | **482.682** | **16355.34** | **78422.23** |  |  |
| 6-HYDROXYFLAVONE | **238.24** |  | **247.7428** | **90.30321** | **8517.499** |  |  |
| WOGONIN | **284.26** |  | **258.2542** | **676.3016** | **2140.941** |  |  |
| BAICALEIN | **270.24** |  | **262.1438** | **65.54845** | **3634.92** |  |  |
| 7,8- DIHYDROXYFLAVONE | **254.24** |  | **277.2542** | **529.6534** | **5820.21** |  |  |
| CHRYSIN | **300.26** |  | **281.4634** | **353.3122** | **916.2911** |  |  |
| APIGENIN | **270.24** |  | **286.2264** | **255.565** | **3634.92** |  |  |
| LUTEOLIN | **286.24** |  | **287.171** | **0.866761** | **1961.631** |  |  |
| DIOSMETIN | **300.26** |  | **296.86** | **11.56** | **916.2911** |  |  |
| TRICIN | **330.29** |  | **355.836** | **652.5981** | **0.057744** |  |  |
| NOBILETIN | **402.39** |  | **414.39** | **144** | **5163.816** |  |  |
| TANGERITIN | **372.37** |  | **493.7092** | **14723.2** | **1750.56** |  |  |
| AZALEATIN | **316.26** |  | **229.8502** | **7466.654** | **203.6415** |  |  |
| GOSSYPETIN | **318.23** |  | **255.776** | **3900.502** | **151.2974** |  |  |
| FISETIN | **286.24** |  | **275.732** | **110.4181** | **1961.631** |  |  |
| QUERCETIN | **302.24** |  | **279.997** | **494.751** | **800.3411** |  |  |
| KAEMPFEROL | **286.24** |  | **282.4686** | **14.22346** | **1961.631** |  |  |
| MORIN | **302.24** |  | **284.052** | **330.8033** | **800.3411** |  |  |
| FUSTIN | **288.25** |  | **285.0612** | **10.16845** | **1787.624** |  |  |
| KAEMPFERIDE | **300.26** |  | **293.442** | **46.48512** | **916.2911** |  |  |
| GALANGIN | **270.24** |  | **299.3532** | **847.5784** | **3634.92** |  |  |
| ISORHAMNETIN | **316.26** |  | **304.4188** | **140.214** | **203.6415** |  |  |
| RHAMNAZIN | **330.29** |  | **310.2051** | **403.4032** | **0.057744** |  |  |
| MYRICETIN | **318.23** |  | **311.7215** | **42.36057** | **151.2974** |  |  |
| RHAMNETIN | **316.26** |  | **327.6834** | **130.4941** | **203.6415** |  |  |
| MYRICITRIN | **464.4** |  | **473.8621** | **89.53134** | **17921.1** |  |  |
| RUTIN | **610.52** |  | **576.1325** | **1182.5** | **78394.23** |  |  |
| DAIDZEIN | **254.24** |  | **305.5832** | **2636.124** | **5820.21** |  |  |
| GENISTEIN | **270.24** |  | **307.2475** | **1369.555** | **3634.92** |  |  |
| CALYCOSIN | **284.27** |  | **318.2088** | **1151.842** | **2140.015** |  |  |
| BIOCHANIN | **284.26** |  | **321.4872** | **1385.864** | **2140.941** |  |  |
| PUERARIN | **416.38** |  | **451.845** | **1257.766** | **7370.171** |  |  |
| EPICATECHIN | **290.27** |  | **326.75** | **1330.79** | **1620.892** |  |  |
| CATECHIN | **290.27** |  | **328.7168** | **1478.156** | **1620.892** |  |  |
| THEAFLAVIN | **564.5** |  | **334.656** | **52828.26** | **54741.82** |  |  |
| EIPGALLOCATECHIN | **306.27** |  | **337.7274** | **989.568** | **588.5622** |  |  |
| EGCG | **458.4** |  | **478.6908** | **411.7166** | **16350.66** |  |  |
| PHLORETIN | **274.26** |  | **313.1416** | **1511.779** | **3166.347** |  |  |
| GARBANZOL | **272.25** |  | **286.796** | **211.5861** | **3396.593** |  |  |
| AROMADEDRIN | **288.25** |  | **293.0886** | **23.41205** | **1787.624** |  |  |
| TAXIFOLIN | **304.25** |  | **302.7016** | **2.397543** | **690.6542** |  |  |
| DIHYDROGOSSYPETIN | **320.25** |  | **318.2959** | **3.818507** | **105.6846** |  |  |
| AMPELOPSIN | **320.25** |  | **332.82** | **158.0049** | **105.6846** |  |  |
| DIHYDROMORIN | **304.25** |  | **354.9824** | **2573.776** | **690.6542** |  |  |
| SILIBILIN | **482.4** |  | **397.5578** | **7198.199** | **23064.41** |  |  |

Supplementary Figure2: Cartoon representation of Parkin protein with the ligands (middle) (A), and interactions between Parkin and the ligands (left) (B) showing the interacting amino acid residues of Parkin and ligplot shows the hydrogen bonding pattern of the ligand with the Parkin protein (C) (presented in the PowerPoint marked as flavonoids).
